# Human ESCRT-I and ALIX function as scaffolding helical filaments *in vivo*

**DOI:** 10.1101/2024.05.01.592080

**Authors:** Stephanie J. Spada, Kevin M. Rose, Paola Sette, Sarah K. O’Connor, Vincent Dussupt, V. Siddartha Yerramilli, Kunio Nagashima, Virginie Helle Sjoelund, Phillip Cruz, Juraj Kabat, Sundar Ganesan, Margery Smelkinson, Aleksandra Nita-Lazar, Forrest Hoyt, Suzanne Scarlata, Vanessa Hirsch, Sonja M. Best, Michael E. Grigg, Fadila Bouamr

**Affiliations:** Laboratory of Parasitic Diseases and Laboratory of Molecular Microbiology, NIAID, NIH, Bethesda, MD, 20894; Department of physiology and biophysics, Stony Brook University, Stony Brook, NY, 11794-8661; Frederick National Laboratory for Cancer Research, Frederick, MD 21701; University of Oklahoma Health Sciences Center, NIAID, NIH, Bethesda, MD, 20894; Computational Biology Section Bioinformatics and Computational Biosciences Branch, NIAID, NIH, Bethesda, MD, 20894; Biological Imaging Section, NIAID, NIH, Bethesda, MD, 2089; Laboratory of Systems Biology, NIAID, NIH, Bethesda, MD, 2089; Department of Chemistry and Biochemistry, Worcester Polytechnic Institute, Worcester, MA 01609; Research Technologies Branch, Rocky Mountain Laboratories, NIAID, NIH, Hamilton, MT, 59840, USA; Laboratory of Neurological Infections and Immunity, Rocky Mountain Laboratories, NIAID, NIH, Hamilton, MT, 59840, USA; Laboratory of Parasitic Diseases, NIAID, NIH, Bethesda, MD, 20894

## Abstract

The Endosomal Sorting Complex Required for Transport (ESCRT) is an evolutionarily conserved machinery that performs reverse-topology membrane scission in cells universally required from cytokinesis to budding of enveloped viruses. Upstream acting ESCRT-I and ALIX control these events and link recruitment of viral and cellular partners to late-acting ESCRT-III CHMP4 through incompletely understood mechanisms. Using structure-function analyses combined with super-resolution imaging, we show that ESCRT-I and ALIX function as distinct helical filaments *in vivo*. Together, they are essential for optimal structural scaffolding of HIV-1 nascent virions, the retention of viral and human genomes through defined functional interfaces, and recruitment of CHMP4 that itself assembles into corkscrew-like filaments intertwined with ESCRT-I or ALIX helices. Disruption of filament assembly or their conformationally clustered RNA binding interfaces in human cells impaired membrane abscission, resulted in major structural instability and leaked nucleic acid from nascent virions and nuclear envelopes. Thus, ESCRT-I and ALIX function as helical filaments *in vivo* and serve as both nucleic acid-dependent structural scaffolds as well as ESCRT-III assembly templates.

**Significance statement:** When cellular membranes are dissolved or breached, ESCRT is rapidly deployed to repair membranes to restore the integrity of intracellular compartments. Membrane sealing is ensured by ESCRT-III filaments assembled on the inner face of membrane; a mechanism termed inverse topology membrane scission. This mechanism, initiated by ESCRT-I and ALIX, is universally necessary for cytokinesis, wound repair, budding of enveloped viruses, and more. We show ESCRT-I and ALIX individually oligomerize into helical filaments that cluster newly discovered nucleic acid-binding interfaces and scaffold-in genomes within nascent virions and nuclear envelopes. These oligomers additionally appear to serve as ideal templates for ESCRT-III polymerization, as helical filaments of CHMP4B were found intertwined ESCRT-I or ALIX filaments *in vivo*. Similarly, corkscrew-like filaments of ALIX are also interwoven with ESCRT-I, supporting a model of inverse topology membrane scission that is synergistically reinforced by inward double filament scaffolding.

## Introduction

Endosomal Sorting Complex Required for Transport (ESCRT) is a heteromultimeric cellular machinery that processes a unique dynamic membrane fission event, termed inverse or reverse topology. This event takes place on the cytoplasmic face of the membrane and is universally required to complete critical cellular processing including cytokinesis, autophagosome closure, and nuclear envelope reformation, and is usurped by enveloped viruses to complete budding, including HIV-1 [1–3]. In the cell, ESCRT-I can interact with ESCRT-III through ESCRT-II, in a set of sequential interactions that involve ESCRT-I VPS28 binding ESCRT-II EAP45, while ESCRT-II EAP-20 binds ESCRT-III CHMP6 [4]. In the case of HIV-1 budding, ESCRT-I interactions with ESCRT-III through ESCRT-II have been recently suggested [5, 6], although the nature of the interactions involved, and their functional significance are debated. A second pathway to link cellular and viral proteins to ESCRT-III CHMP proteins is mediated through ALIX, which binds ESCRT-III directly and constitutes an independently functional pathway in both cellular [7] and viral [8] contexts.

The first indication that the ESCRT-I complex may form higher order oligomers came from the Williams’ laboratory structure of the yeast ESCRT-I core [9] (PDB code: 2CAZ) which observed that the yeast ESCRT-I can assemble into helical filaments **(also see supplemental Figure 1)**. These filaments consisted of 12 ESCRT-I cores per turn oriented at a 9° helical angle, properties that are similar to the recently resolved structure of an ESCRT-III helical polymer [10]. Remarkably, an average of a dozen ESCRT-I molecules are estimated to form in budding necks [11, 12], suggesting only a subset of Gag molecules recruit ESCRT-I through TSG101. Additionally, similar filaments of human ESCRT-I were recently captured in solution as well as via crystallography [13]. We hypothesized that the predicted filament structure is likely biologically relevant as a template for the oligomerization of the human ESCRT-I complex *in vivo*. These observations also suggested that ESCRT-I as well as ALIX, might form these structures in cells to recruit and might serve as a template for ESCRT-III filament assembly, thus bypassing ESCRT-II, as studies with HIV-1 previously reported [14].

Both ALIX -or Bro1-containing proteins and ESCRT-I TSG101 are recruited to membrane abscission sites of viral budding necks (HIV-1 [15, 16]). Dual recruitment has also been detected in organelles where pathogens are surviving and replicating, at pathogen invasion sites and areas of resource uptake (as in the case of toxoplasma gondii) [17, 18], cytokinetic bridges [19], multivesicular bodies [20], and sites of plasma membrane and endosomal membrane repair [21, 22]. Recruitment of both ESCRT-I and ALIX or a Bro1 domain containing ortholog of ALIX, where both are necessary to process membrane scission, also includes sites of neuronal pruning in Drosophila melanogaster [23], and cytokinesis abscission sites [24]. Additionally, both TSG101 and ALIX are required for the recruitment of ESCRT-III to cytokinetic bridges in *Drosophila melanogaster* and *C. elegans* [25]. Dual recruitment of ESCRT-I and ALIX hints to a key synergistical role for ALIX/ALIX-like proteins and ESCRT-I co-association at sites of inverse topology membrane scission.

ESCRT-I and ALIX are recruited to most if not all sites of inverse topology membrane scission pertinent to health and disease. The nature of their roles in live cells as obligatory adaptors that link cellular and viral proteins to ESCRT-III, has remained elusive until recently [13]. Here we used biochemical and functional assays, live and super-resolution imaging along with modelization derived from known structures to independently elucidate underlying mechanisms. Using HIV-1 budding and cytokinesis as models, we found that ESCRT-I and ALIX polymerize into helical filaments that assemble along the lumen of membrane abscission necks *in vivo*, and are dependent upon the evolutionarily conserved interfaces in ESCRT-I VPS28 and the ALIX Bro1 domain, respectively. When assembly of such filaments is disrupted, nascent virions cannot be released and dividing daughter cells exhibited loss of nucleic acid compartmentalization and structural instability, indicating ESCRT-I and ALIX filaments are critical structural scaffolds during viral and cellular membrane scission. Here we show these functions require newly clustered functional interfaces; the first binds negatively charged membrane (or RNA), while the second assembles ESCRT-I and ALIX subunits into helical filaments. ESCRT-III CHMP4B helical filaments were also found interlaced with those of ESCRT-I or ALIX in lumens of abscission bridges *in vivo*. Both ALIX and ESCRT-I VPS28 filaments also intercrossed therein, supporting a synergistical model. We show that ALIX and ESCRT-I subunits assembling into helical filaments is necessary for optimal structural scaffolding and genome compartmentalization *in vivo* while simultaneously forming templates for ESCRT-III polymerization, both key to structural stability and inverse topology membrane scission. This study addresses a gap in the sequential progression that leads to inverse topology membrane scission and enhances our understanding of how the early binding adaptors ESCRT-I and ALIX might mechanistically initiate ESCRT-III-mediated membrane sealing.

## Results

### Budding defects result in accumulation of structurally unstable HIV particles at the plasma membrane

Recruitment of both ESCRT-I subunit TSG101 and ALIX to bind ESCRT-III [26, 27] has often been observed during viral (HIV-1 [15] and Ebola virus [28]) and cellular membrane scission (cytokinesis: ALIX and TSG101 [19]; neuronal pruning: HD-PTP and TSG101 [23]). While ESCRT-III was shown to polymerize into filaments at sites of membrane budding [29], how ESCRT-I or ALIX trigger the assembly of ESCRT-III *in vivo* remains unknown. We used HIV-1 budding necks that recruit both ALIX and ESCRT-I TSG101 through the canonical binding L domains YPXnL and PTAP, located in HIV-1 Gag p6 domain, as a model to examine membrane scission sites and elucidate mechanisms of downstream recruitment.

Mutations in the L domain PTAP and YPXnL are known to cause disruption of the viral structural protein Gag maturation by the viral Protease (Pr), a hallmark of budding defects, phenotypes recently explained by Gag-Pol polyprotein gradual seepage out of particles, once Pr is activated in L domain-devoid-developing virions [30]. Consequently, unprocessed Gag products accumulate and are detected in nascent virions. We used the retention of Gag-Pol (Pr) [31] and in turn inefficient Gag maturation -- to evaluate the extent of budding defects, once ESCRT-I TSG101 or ALIX binding L domains were disrupted. Total cell lysates and isolated flotation fractions [32, 33] (**Figure 1a** and **Figure 1b** upper), where Gag and maturation products accumulated at the PM, were assessed for expression and accumulation of processed viral proteins. Western blot analysis revealed that both anti-NC (**Figure 1b**) and anti-p6 antibodies (**Figure 1a**) recognize the full length Gag, as well as maturation product NC-p6 at the C-terminal end of the protein, while an anti-MA antibody completes the virus maturation panel of Gag at these sites of virus assembly and budding (**Figure 1b, lower)**. WT HIV-1 showed no accumulation of unprocessed structural Gag proteins or other maturation defects and both Gag and its NC-containing maturation product are detected at the PM, (**Figure 1b**, right and inset with longer exposure). In contrast, we observed severe maturation defects with the HIV-1 PTAP-mutant (**Figure 1b**, lane 2 and inset with longer exposure-right). NC mature protein was diminished or lost and a clear and reproducible diminution of p7NC cleavage from Gag was observed as evidenced by a consistent accumulation of immature p15NCp1-p6 (**Figure 1b**, lane 5). An intermediate phenotype was observed with HIV-1 YPXnL-mutant, where Gag NC- and p6-maturation intermediates accumulated in cell extracts probed with either anti-p6 (**Figure 1a** left lane 1) or anti-NC antibodies (**Figure 1b**, left lane 3). Similar defects were observed at the PM in floatation fractions (**Figure 1b**, right, lane 6), where unprocessed p15NCp1-p6 also consistently accumulated. Both PTAP- and YPXnL-mutants displayed a decrease in NC maturation efficiency. Since both mutants are known to fully assemble stable nascent virions, we interpreted the maturation defects observed as likely a premature loss of Pr (Gag-Pol) from nascent virions at the PM (**Figure 1a** and **1b** left, lane 3 and longer exposures in **insets**). These defects were further confirmed with an anti-MA antibody that also detected the accumulation of unprocessed Gag maturation intermediates (**Figure 1b** lower, lanes 5 and 6, marked with red asteriks). Maturation defects of PTAP- and YPXnL-mutants were reproducible and evaluated to be of ∼70% and ∼15%, respectively, in six independent experiments **(Figure 1e**). These defects indicate that viral structural integrity is compromised and there is a lack of retention of all components in developing virions at sites of membrane closure.

**Figure 1:**
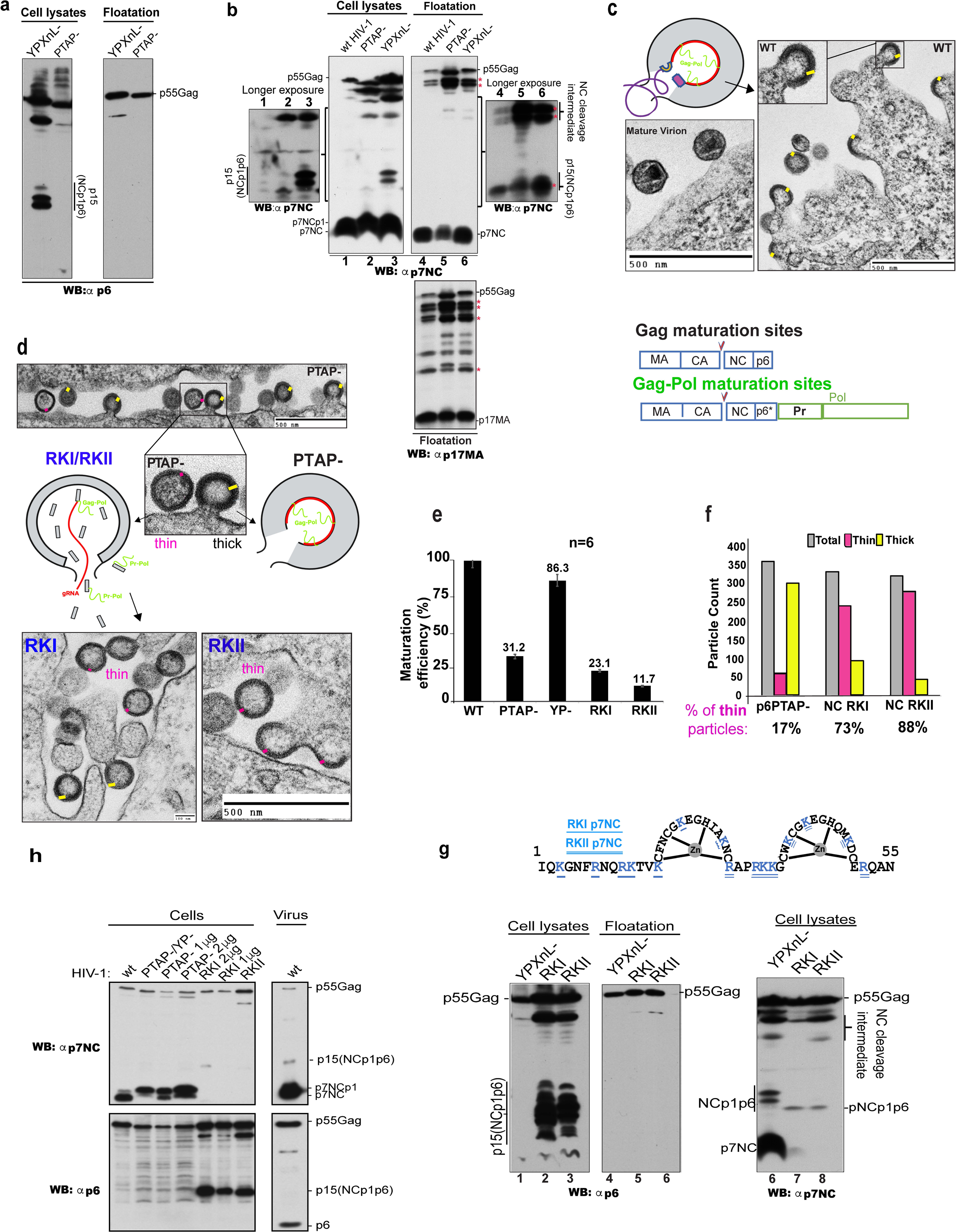
Preventing recruitment of upstream ESCRT-I and ALIX cause structural instability in developing enveloped virions at the PM. (**a**) and (**b**) Examination of HIV-1’s scission of viral budding necks at PM using floatation assays. Cell lysates from WT HIV-1, YPXnL- or PTAP-mutant were loaded on floatation gradients and their protein content subsequently analyzed by SDS-PAGE and western blotting using antibodies to NC (**b**) upper panel, against MA (lower panel) or p6 (**a**). Accumulated unprocessed Gag maturation products are marked with red asterisks in darker exposure insets from portions of membranes marked with side accolades are shown next to western blot panels in (**b**). (**c**) A schematic representation of a WT HIV-1 budding particle that recruits ESCRT-I (yellow), ALIX (purple) and ESCRT-III filaments that assemble in the lumen of viral budding necks (dark purple) to release mature virions with conical internal cores. Conversely, HIV-1 lacking ESCRT-I and/or ALIX remain open at the PM (**d**), with an activated viral Pr. Gag/Gag-Pol (blue and green boxes) are first cleaved between CA and NC (arrowheads) and NC containing Gag/Gag-Pol maturation products (green), associated with gRNA (red) are gradually leaked from arrested particles. RKI and RKII thin virions are shown in electron micrographs as lowest panels. **(e)** Loss of Pr led to accumulation of unprocessed Gag products (detailed in b and h), which are used to evaluate Pr maturation inefficiency in 6 independent experiments. (**f)** Lattice leaks or defects are visible by TEM as discrete morphological changes observed with PTAP-mutant and ∼17% of particles exhibited thinner shells (pink bars), consistent with the loss of Gag/Gag-Pol p15NCp1-p6 lattice, in contrast to WT Gag thick shells (yellow bars) where the gRNA dark electron dense lattice is observed. Thin particles of RKI and RKII mutants exhibited ∼73% and ∼88%, respectively, structurally instable thin, budding virions, in two separate experiments and more than 360 virions counted/sample. (**g**) Schematic depiction of HIV-1 RKI and RKII constructs used in this study. (**g)** Data from experiments like those conducted in (**a**) and (**b**) are shown, with RKI and RKII mutants compared to WT, PTAP-. Cellular extracts from mutants RKI and RKII are analyzed by western blot with anti-NC (upper) and anti-p6 (lower) antibodies, and their maturation patterns compared to those of PTAP-viruses, and double mutant (PTAP- and YP-) virus, before examination of nascent virions maturation and protein content using flotations assays (**h**). Loss of p15NCp1p6 maturation product from RKI and RKII is noted and p15NCp1p6 accumulates in cells (h, left) but is absent in developing virions at the membrane (h, right), in 6 independent experiments (like data in **g**). The absence of NC maturation product, that also derives from p15NCp1p6 processing by Pr, was not detected with an anti-NC antibody (**h**, third panel right), despite the antibody readily recognizing WT and mutants RKI and RKII unprocessed Gag proteins equally, as can be seen in both **h** and **g** panels.

Maturation defects are consistent with the loss or seepage of NC-containing Gag and Gag-Pol polypeptides [30], both of which associate with viral RNA genomes [34] through interaction first with NC and then the Gag-Pol encoded Integrase, also a known constituent of the viral ribo-nucloprotein (RNP) [35]. Transmission electron microscopy (TEM) analysis of developing virions of the PTAP-mutant (in six independent experiments) showed that most virions exhibited immature particles morphologically indistinguishable from WT virions and harbor a dark genome-binding GagNC-p6 layer -the RNA-p15NCp1p6 density [36], referred to below as “thick” particles. Conversely, ∼ up to 20% of a second group showed “thin” particles where no dark genome-binding GagNC-p6 layer was observed (**Figure 1d, pink**). Consistently, no “thin” particles were observed in the WT virions sample. Similar phenotypes were previously described by Carlson and colleagues [36] with L domain mutants as novel Gag lattice intermediates that retained only densities corresponding to MA-CA assemblies but lacked the NC-gRNA-p6 layer, in agreement with loss of NC-gRNA assemblies (NCp1p6-RNPs) we observed in PTAP-mutant thin particles (**Figure 1d** and quantified in **1f** ). A clear and reproducible reduction of virus maturation was also observed with the YPXnL-mutant virus (**Figure 1e**), a finding consistent with a structural role for ALIX in stabilizing developing virions at the PM and in turn retention of Gag-Pol therein (Figure **1a and b**). Collectively, the data here support a model in which ESCRT-I and ALIX provide key structural scaffolding to assembled developing virions prior to their budding and release by ESCRT-III polymers [11].

### Nascent virions that incorporate RNA genome are favored for ESCRT-dependent abscission

Next, we sought to decipher the mechanisms that govern ESCRT-I and ALIX scaffolding and the retention or stabilization of NC-associated viral gRNA within nascent virions at the PM. Viral particles released in the presence of either cellular or gRNA showed seemingly indistinguishable morphologies, initially suggesting that gRNA is not necessary for virus assembly and budding [37]. Recent studies however showed that the nature of RNA incorporated during virus assembly is important, as longer RNA [38] (also reviewed in [39]) and the presence of Psi pacakaging signal enhanced and accelerated virus release [40–43], suggesting viral genomes might be involved in efficient virus budding and abscission [44]. We used Gag mutant viruses, RKI and RKII that lost the ability to incorporate gRNA but initiated assembly of nascent virions at the PM [45] **(Figure 1h**), to assess the role of gRNA in virus budding and abscission. Although both mutant RKI and RKII Gag proteins associated with the PM in floatation assays (**Figure 1h, lanes 5 and 6**), neither mutant Gags fully cleaved NC-p6 intermediate maturation product into NC as evidenced by p15 accumulation detected by the anti-p6 antibody (**panel 1h, lanes 2 and 3**) and the absence of fully mature NC (panel **1h, lanes 2, 3, 7 and 8**). Severe Gag processing defects were quantified from maturation panels (**Figure 1g left and 1h**) to be of ∼ 77% and 89% for RKI and RKII, respectively, (**Figure 1e)** and were observed in genome-devoid nascent virions, as high amounts of immature Gag intermediates accumulated. Such a loss was also detected in TEM micrographs of RKI and RKII virus mutants (**Figure 1d**). Maturation defects were more severe that those seen with the PTAP-mutant (only ∼20% of thin virions) as ∼ 73% and 88% of RKI and RKII developing virions, respectively were found to form thinner Gag-shells virions (**Figure 1f**), findings consistent with loss of p15 NCp1-p6 layer at the PM (**Figure1h**) and aligned with quantification of maturation defects shown in **Figure 1e**. The absence of NC or NC-containing polypeptides is not due to loss of recognition by the anti-NC antibody, as both RKI and RKII Gag proteins were detected comparably to WT Gag (**panels 1g and 1h**). As RNA genomes are also important for virus assembly, we concluded that despite the accumulation of seemingly assembled spherical particles, these nascent virions displayed a combination of assembly and budding defects that could be attributed to a lack of proper virion stabilization and scaffolding by ESCRT-I and ALIX at the membrane.

Next, we assessed if gRNA incorporating signal Psi, is important for gRNA’s role in ESCRT-I and ALIX structural scaffolding within developing virions at the membrane. We examined whether providing a fully functional (Psi+) gRNA-binding construct *in trans* in virus rescue assay could restore ESCRT scaffolding functions (i.e.: RK mutants). RKII nascent virions (and RKI, *not shown*) failed to process Gag fully or retain NC-gRNPs within nascent virions at the PM as assessed by an anti-NC antibody (**Figure 2b**, lane 1). Restoring virus maturation and structural integrity to these mutant viruses was attempted with two release-defective HIV-1 constructs (**Figure 2a)**. The first construct was HIV-1 Psi (double L Domain Mutant) DM, which carries a WT Psi sequence and two inactive L domains (PTAP-/YP-); the second construct encodes for a Gag DM that lacks Psi but carries a functional NC within Gag expressed under a CMV promoter. Co-expression *in trans* (∼ 5%) of HIV-1 Psi DM with RKII restored robust virus release in a dose dependent manner (**Figure 2b**, lanes 2-4), in contrast to Gag DM (no Psi), which performed far less efficiently and only at ∼40% of the WT levels (+Psi) (**Figure 2b, lanes 5-7**). Virus stabilization and release was however increased when co-expression of Gag DM with RKII NC mutant was incrementally increased as previously reported [40], albeit at a much diminished efficiency (∼ 40%) compared to that observed with Psi carrying constructs, in three independent experiments (**Figure 2b, and** quantification graph below). Furthermore, increasing amounts of an HIV-1 DM lacking gRNA-incorporating Psi loops S1 and S3 [46], reproducibly displayed a marked reduction, down to ∼ 39% of virus budding (**Figure 2c**, and quantification graph below). This data indicates that gRNA incorporation is critical for efficient scaffolding and stabilization of developing virions by ESCRT-I and ALIX and underscores the pivotal role of Psi-containing gRNA in accelerating the rate of HIV-1 budding.

**Figure 2:**
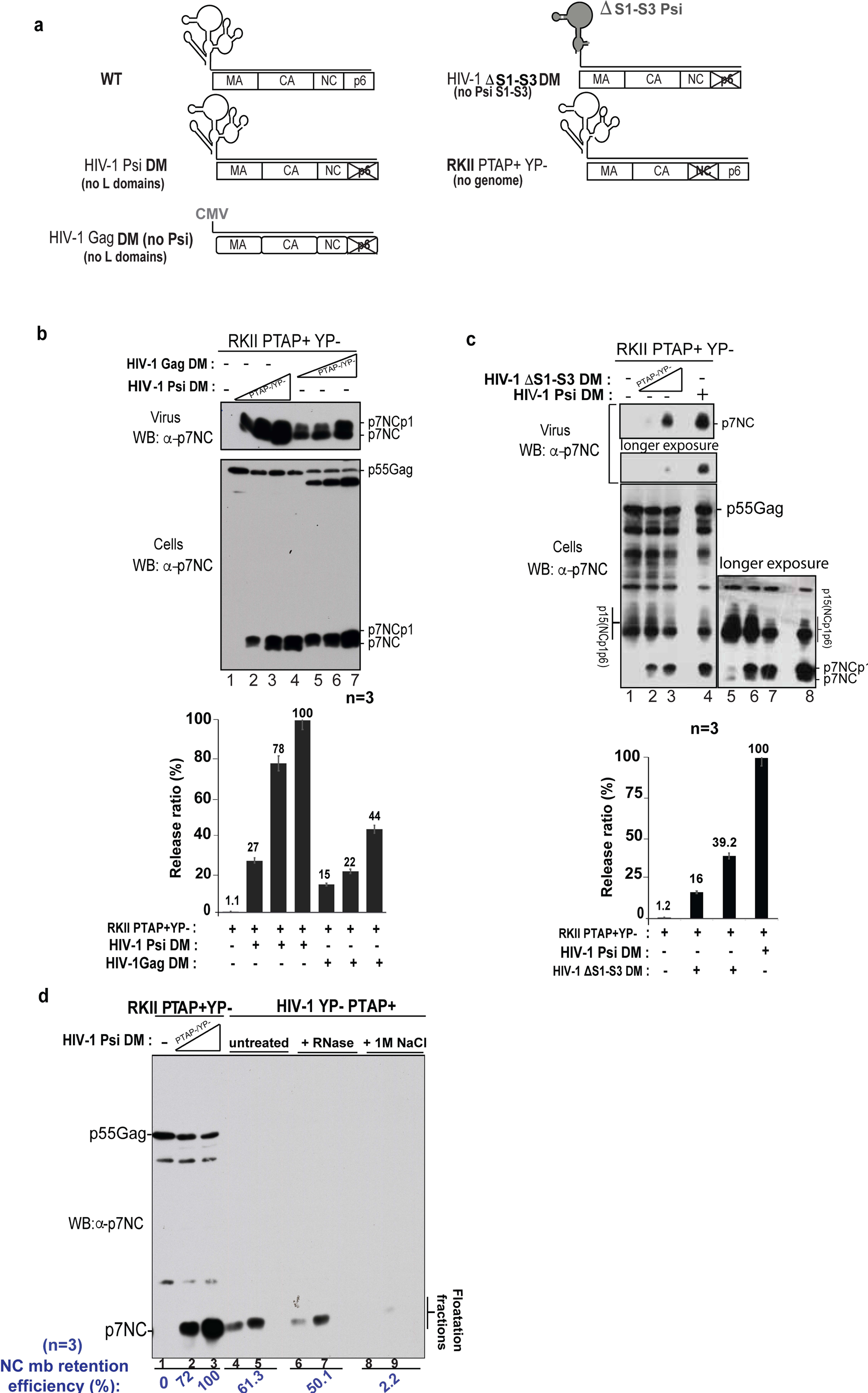
gRNA Role of nucleic-acid genomes in ESCRT-dependent virus abscission. (**a**) Schematic representation of constructs containing a competent virus (WT), missing the PTAP and YPXnL L domains (HIV-1 Psi DM), lacking only RNA-binding residues in NC domain (RKII PTAP+ YP-), missing the entire 5’ UTR region and PTAP and YPXnL L domains named HIV-1 Gag DM (no Psi) or a construct carrying a Psi-mutant region and lacking PTAP and YPXnL L domains (HIV-1 deltaS1-S3 DM) were used. (**b**) Virion release defective RKII PTAP+ (lane 1) was robustly rescued with increasing amounts of HIV-1 Gag Psi DM virus (lanes 2-4; ∼ 100%) and with a less efficiency by the HIV-1 Gag DM (lanes 5-7; ∼40%), evaluated from three experiments in the quantification graph below blot. (**c**) Similar Virus rescue experiments as in **b** have been performed to rescue the RKII PTAP+ mutant (lane 1); Virus release was robustly rescued by HIV-1 Psi DM (lane 4), while the mutant HIV-1 8S1-S3 (loops) DM functions only at ∼39% in three independent experiments as shown in the quantification graph below blot. (**d**) Virus rescue experiments as in **b** were performed to assess if RKII PTAP+ YP-virus rescue can be captured at the PM using floatation assays. RKII PTAP+ YP-showed no mature NC protein in flotation assays at the PM (lane 1), while co-expression of increasing amounts of HIV-1 Psi DM restored virus stability and retained NC-containing and mature NC protein(s) (lanes 2 and 3). Treatment of fractions with RNase (lanes 6 and 7) or high salt at 1M NaCl detached NC complexes (lanes 8 and 9), as assessed in three independent experiments; membrane attachment efficiency values are listed below the western blot with the anti-NC antibody.

This hypothesis was further tested by providing *in trans* ∼10% of HIV-1 Psi DM construct -that binds gRNA but lack ESCRT binding- to the RKII mutant virus, which binds ESCRT but lack gRNA, and checked stability of nascent virions at the PM using flotation assays. Supplementing HIV-1 Psi (+) DM *in trans* robustly restored virus maturation and the efficient cleavage of NC of RKII, which was observed at the PM in developing virions in floated fractions, and only in response to incremental increasing amounts of HIV-1 Psi (+) DM (**Figure 2d, lanes 1-3**). This result indicated that gRNA restored virion structural stability as well as proper maturation through Gag-Pol retention, suggesting nascent virions regained use of ESCRT scaffolding which supports a functional link between gRNA and ESCRT function at the PM. Importantly, upon treatment with RNase, these nascent virions therein were minimally affected (**Figure 2d**, compare **lanes 6 and 7),** with comparable association with the PM **(lanes 4 and 5**) indicating that the gRNA genome is shielded away from the enzymatic effects of RNase. Conversely, treatment with 1M NaCl detached NC-associated gRNA from nascent virions (**lanes 8-9)** and their association to the PM plummeted to ∼2.2%. This result further supported the view that gRNA is shielded within nascent virions, while the integrity of all structural components held together as a macrostructure within the developing virion is sensitive to salt treatment. Considered collectively, this data suggests that nascent virions that assemble around gRNA exhibit the highest structural integrity and stability by recruiting and cementing all structural and enzymatic components together to complete viral maturation through a an ESCRT scaffolding-dependent mechanism.

### ESCRT-I assembles into helical filaments critical for inverse topology membrane scission

Although ESCRT-I is recruited during Gag assembly, virion-associated TSG101 becomes “visible” only when virus assembly is complete and budding of nascent virions initiated [11], suggesting Gag assembly is important and necessary for ESCRT-I concentration. Our data showed that developing virions that incorporate gRNA displayed the most stable particles, and efficiently retained their gRNPs (**Figure 2**), suggesting that ESCRT-I stabilization or scaffolding of nascent virions involved accumulation of ESCRT-I structures therein [47]. Indeed, a subcomplex of ESCRT-I TSG101-VPS28 was recently assembled *in vitro* [13]. We asked whether similar structures are formed by the entire ESCRT-I complex *in vivo* to maintain virus stability at the PM (**Figure 2**), a view consistent with the observation that ∼ 95% of HIV-1 released harbor RNA genomes [44]. Structurally, ESCRT-I is organized as a core composed of three parallel helical hairpins belonging to VPS28, TSG101 (yeast VPS23), and VPS37 [48, 49] with an elongated stalk and flexible tethers extending away from the core. The latter are believed to capture ESCRT-I viral and cellular partners such as ubiquitinated proteins, lipids and downstream acting ESCRT complexes [9, 48, 50]. The Williams’ laboratory structure of the yeast ESCRT-I core [9] (PDB code: 2CAZ) first showed that ESCRT-I can form high order oligomers, or helical filaments **(Supplemental Figure 1, panels a and b)**. The filaments consist of 12 ESCRT-I cores per turn oriented at a 9° helical angle, properties that are similar to the recently resolved structure of an ESCRT-III helical polymer [10]. Remarkably, an average of a dozen ESCRT-I molecules are estimated to coat budding necks [11, 12], suggesting only a subset of Gag molecules is engaged in recruiting ESCRT-I through TSG101. Similar filaments of human ESCRT-I were observed in solution as well as via crystallography [13], suggesting this filament structure is likely biologically relevant.

Based on our previous experiments and known structures of ESCRT-I and HIV proteins, we generated a structural model of a virus budding neck scaffolded by ESCRT-I (**Figure 3a).** ESCRT-I copolymers are dependent upon VPS28-mediated oligomerization between adjacent ESCRT-I cores via an evolutionarily conserved interface in VPS28 (**Figure 3d).** In support of this model, abrogation of this interface in VPS28 by mutagenesis abolished proper assembly of ESCRT-I polymers in cells (**Figure 3e)**. Given the known parallels between virus budding and cytokinesis, we next asked whether or not ESCRT-I scaffolding was important for cytokinetic abscission. To test this, we used STED super-resolution fluorescent imaging to probe for the assembly of ESCRT-I scaffolds in cytokinetic bridges between two dividing daughter cells. These sites sites are known to use ESCRT-I to complete inverse topology membrane scission in an ESCRT-III dependent manner, and resembles the process used for scission scission of HIV-1 budding necks as well. We observed ESCRT-I helical filaments along abscission bridges in HeLa cells expressing mCherry VPS28 fusion proteins. Corkscrew-like filaments decorated the entire inner side of these bridges, a representative example is shown in **Figure 3b**. This data showed that ESCRT-I assembles *in vivo* into helical filaments, in agreement with our model in **Figure 3a**. ESCRT-I helical filaments exhibited gradual diminished diameters when the membrane constricted near the cleavage midbody. Measurements reveal filament diameters to range from approximately 60 nm to ∼ 1 um (**Figure 3c and also see details in Suppelemental Figure 3c).** These sizes correspond to established sizes HIV-1 budding necks, estimated to measure ∼50 to 90 nm [51] , [52], [53]- as well as cytokinesis abscission bridges whose size is estimated to range from ∼ 200 nm to 1 um in diameter [29] and also reviewed in [54]).

**Figure 3:**
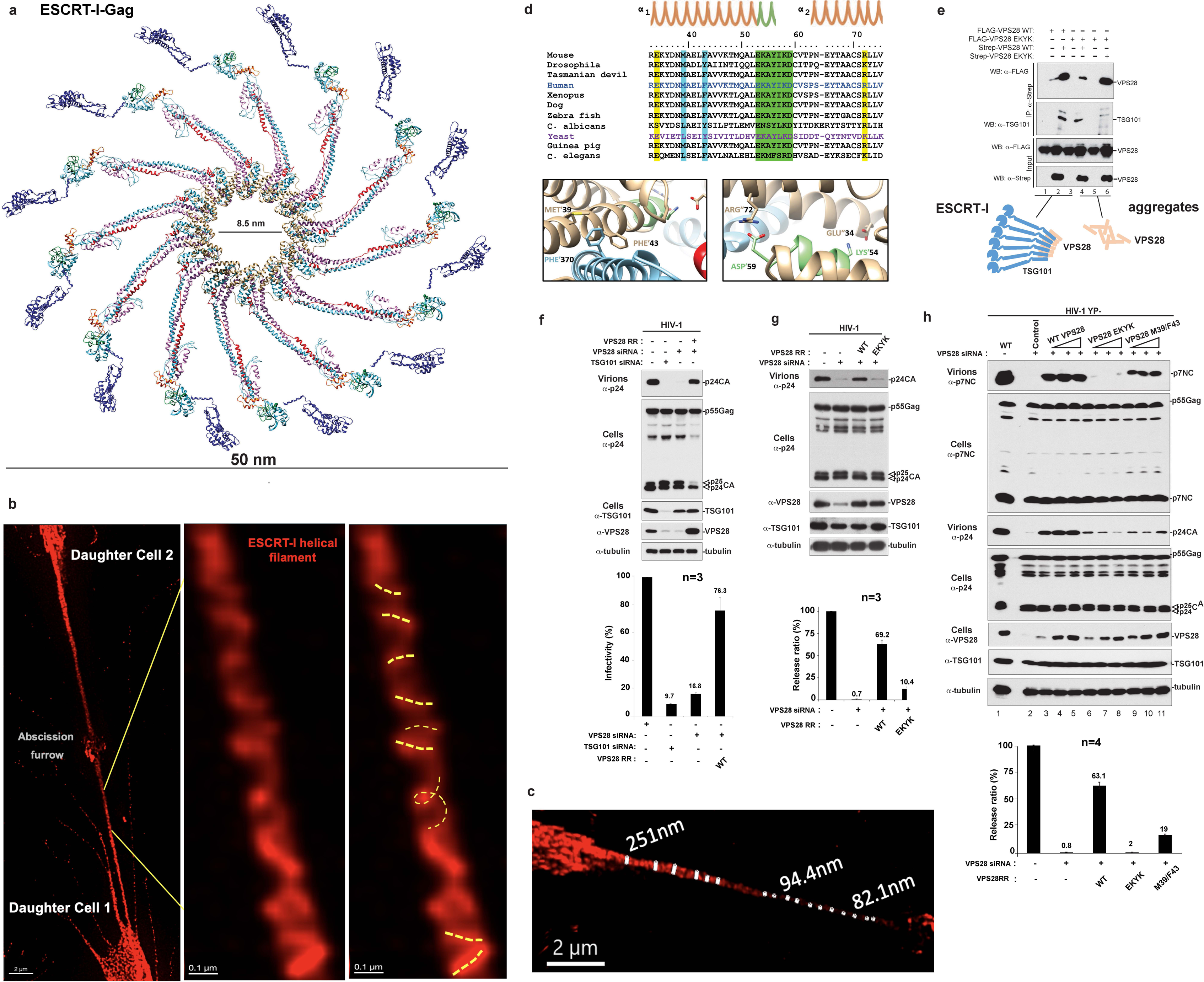
ESCRT-I function requires a new VPS28 interface critical for virion abscission. **(a)** Molecular model of a human ESCRT-I filament engaged with the PTAP sequences of HIV Gag based on the yeast filament structure. **(b)** ESCRT-I helical filaments assemble *in vivo*: Super-resolution imaging was used to capture ESCRT-I filaments in dividing HeLa cells. VPS28-mcherry fusion protein was expressed to track ESCRT-I within abscission bridges between two daughter cells, bar: 2um. Helical filaments have been observed within multiple fields and four independent experiments, a representative of these images is shown, and high magnifications zooms of filaments shown in insets, where helical turns are highlighted with yellow discontinued lines (right), bars 0.1uM **(c)** ESCRT-I helical filaments diameter measurements: Imaris measuring tool was used to estimate the diameter of ESCRT-I filaments as they inwardly coat scission bridges, the size ranges from ∼ 250 to 80 nm. **(d)** Sequence alignment of the N-terminal domains (NTDs) of VPS28 reveals high conservation across species, especially at the _53_EKAYIKDC_60_ sequence (EKYK, highlighted in green). Yeast VPS28 forms hydrogen bonds between neighboring VPS28 molecules, via EKYK and conserved residues on the neighboring molecule (highlighted in yellow, residues shown in bold). An additional predicted contact between yeast VPS28 and VPS23 (TSG101 in humans) was identified (residues shown in bold with blue highlight). Modeling of the human ESCRT-I complex reveals that the _53_EKAYIKDC_60_ sequence forms the equivalent contacts as observed in yeast. A zoom (below alignment, left) is shown that focuses on the specific interactions between K54 and D59 of one human VPS28 with E34 and R72 on the adjacent VPS28 molecule, forming bonds respectively. A second zoom (right) from the human ESCRT-I molecular model highlights the VPS28-TSG101 sub-complex interface. Human VPS28 residues M39 and F43 are predicted to form hydrophobic contacts with F370 in human TSG101. **(e)** Co-immunoprecipitation with WT VPS28 and a VPS28 (EKYK) mutant. 293T cells were transfected with FLAG-tagged constructs of either WT VPS28 (lane 1) or the EKYK mutant (lanes 3 and 5), or a combination of FLAG-tagged and Strep-tagged constructs (lane 2 and 4 for Strep-VPS28 WT, lane 6 for Strep-VPS28 EKYK). Cell lysates were collected and subject to anti-Strep beads, and protein expression levels were verified by western blot. **(f)** HIV-1 virus rescue assay upon depletion of endogenous VPS28. RNAi knockdown of endogenous VPS28 and reconstitution with WT VPS28 restores the release of virions by HIV-1 mutants that have the YPXnL L domain abrogated. 293T cells were transfected with control RNAi or RNAi targeting VPS28 but also with mutant HIV-1 proviral DNA (1 to 1.5 μg), and either empty pcDNA or a version expressing RNAi Resistant (RR) WT VPS28. Intracellular expression of VPS28 and TSG101 was verified when either protein was depleted, as shown in a representative example of three independent experiments-value release efficiency graph below blot. **(g)** Assessing the role of VPS28 EKYK sequence in ESCRT-I mediated HIV-1 release. Similar experiments as in (f) were performed and virus rescue was tested with either WT or mutant VPS28 as indicated above the panel. Virion production and Gag expression levels were analyzed by Western blot with anti-NC and anti-CA antibodies. Tubulin, VPS28, or TSG101 were detected with a mouse anti-tubulin, a rabbit anti-VPS28 (Sigma), or a mouse anti-TSG101 antibody (Novus Biologicals). The experiment shown is a representative of three independent experiments-value release efficiency graph below the blot. **(h)** Comparing VPS28 WT to EKYK and M39/F43D VPS28 mutants’ function in HIV-1 budding. Experiments were conducted as described above with incrementally increasing amounts of VPS28 WT (lanes 3-5), EKYK mutant (lanes 6-8) or the M39/F43D VPS28 mutant (lanes 9-11). The experiment shown is a representative example of four independent experiments-value release efficiency graph below blot.

Sequence alignment of metazoan VPS28 proteins showed a highly conserved NTD, within the short sequence _53_EKAYIKDC_60_, which retained a near identical sequence from yeast to humans (**Figure 3d**). Human VPS28-VPS28 NTDs showed VPS28 NTD1 _54_K and _59_D residues within interacting distance of VPS28 NTD2 _72_R and _34_E residues, respectively (**Figure 3b, see insets**), identifying _53_EKAYIKDC_60_ sequence as a potential VPS28-VPS28 homo-oligomerization determinant within ESCRT-Iwhere neighboring VPS28 molecules are joined by contacts between _55_K and _60_D in NTD1 with _35_E and _73_K in NTD2. Closer sequence analysis revealed that the binding partners of residues within the _53_EKAYIKDC_60_ sequence were also highly conserved, in agreement with the idea that the_53_EKAYIKDC_60_ sequence may be of evolutionary and functional importance. Immunoprecipitation assays were used to assess this hypothesis (**Figure 3c**) and showed that a streptavidin-tagged (OS)-WT VPS28 immobilized on beads captured WT Flag-tagged VPS28. Conversely, it failed to capture a Flag-VPS28 mutant where residues _54_K and _56_YIKD_59_ were substituted with alanines (termed VPS28 EKYK) (**Figure 3e, compare lanes 2 and 4**). Interestingly, endogenous TSG101 was captured in both cases (**Figure 3e, compare lanes 2 and 4)**, suggesting that a WT VPS28-TSG101 subcomplex associates with additional functional VPS28-TSG101 subunits and can further grow these assemblies, in contrast to the mutant VPS28 EKYK which failed to homo-oligomerize with this assembled complex. Similarly, Flag-VPS28 EKYK mutant self-associated with OS-VPS28 EKYK mutant(s) into aggregates whose ability to bind endogenous TSG101 was abrogated (**Figure 3e, lane 6).** However, these VPS28-VPS28 complexes are non-functional as they failed to recruit TSG101 (**lane 6**). These relationships can be explained by our structural model, where an additional point of contact of TSG101 residue _370_F is predicted to interact hydrophobically with residues _39_M and _43_F of VPS28 (**Figure 3d, inset**). Consistent with this, mutation of this _370_F and neighboring residues in TSG101 (_367_RKQF_370_) to alanine interfered with ESCRT-I function in HIV-1 release [26] **(Figure 3f-h)**. Similarly, substitution of VPS28 _39_M and _43_F to D reduced virus release (see below), confirming the possible additional contribution of these residues to ESCRT-I polymerization. Depletion of cellular VPS28 eliminated HIV-1 production similarly to the adverse effect observed upon depletion of cellular TSG101 (**Figure 3f**). In the absence of endogenous TSG101, the cytoplasmic acumulation of the ESCRT-I VPS28 subunit was diminished (**Figure 3f, lanes 2 and 3**). Similarly, RNAi depletion of VPS28 decreased TSG101 function in cells and diminished virus infectivity to ∼16% in comparison to the control conditions **(Figure 3f, lane 3** and infectivity titrations from three independent experiments), further confirming that TSG101 and VPS28 are functionally interdependent [55]. Reintroduction of VPS28 (RR) (**Figure 3f**, **lane 4**), rescued virus release and maturation as evidenced by near full processing of mature viral capsid (p24CA) and a clear decrease of immature viral capsid (p25CA) in cells. Conversely, HIV-1 release was not recovered by the VPS28 EKYK mutant (**Figure 3g, lane 4, and Figure 3h, compare lanes 3-5 to lanes 6-8).** VPS28 M39/F43D mutations were also deleterious to ESCRT-I function but did not completely obliterate function since ∼ 20 to 30% of virus release was retained (**lanes 9-11**), findings confirmed with both anti-CA and anti-NC antibodies. The results here show that ESCRT-I assembles into helical filaments *in vivo* and uncover the functional importance of their highly conserved self-assembly interface in inverse topology membrane abscission.

### ALIX assembles into helical filaments whose disruption along a new Bro1 interface abrogate membrane abscission

The equine lentivirus EIAV relies on ALIX to release progeny virions, implying ALIX can fulfill similar roles to those uncovered above with ESCRT-I. Evidence supporting such a role for ALIX comes from previous work demonstrating that dimeric ALIX decorates and bridges CHMP4B filaments that have a diameter of ∼50 nm [56]. To further define such a conserved role for ALIX in the lumen of scission sites, the sequences and crystal structures of ALIX and its Bro1 domain containing ortholog Brox were compared. Sequence alignments of ALIX and homologues from several species revealed a highly conserved _317_FIY_319_ motif located at the C-terminal tip of Bro1 in ALIX as well as ALIX homologues Brox and HD-PTP (**Figure 4b**). Structural analysis demonstrated that this motif lies at the center of a multitude of different Bro1-Bro1 interfaces. Most notably, in the presence of Bro1-binding ESCRT-III CHMP4B peptide, Brox adopts an unusual lemniscate wire within the crystal that occasionally intersects as a four-way junction of Bro1 domains with two FIY motifs oriented opposite one another at the center of this apex (PDB code: 3UM3, [57]) (**Supplemental Figure 2**). Additionally, structural modeling revealed that the Brox Bro1 domain alone forms trimeric arrangements that places one FIY motif at each interface (**Supplemental Figure 2**). Additionally, in the presence of Brox Bro1-binding of an ESCRT-III CHMP5 peptide, a pair of Bro1 domains are held together at a dimer interface with two FIY motifs in direct contact with one another (PDB code: 3UM0, [57]) (**Supplemental Figure 2**). Finally, a pair of Bro1 domains in a crystal structure of HD-PTP in complex with CHMP4A are held together by spatially separated FIY motifs (PDB code: 5MK1) (**Supplemental Figure 2**). The biological relevance of several of these new interfaces were supported by Complexation Significance (CSS) scores of 0.8 or better, on a scale of 0 to 1, with 1 highlighting an interface as critical for complexation [58] (**Supplemental Figure 2**).

**Figure 4:**
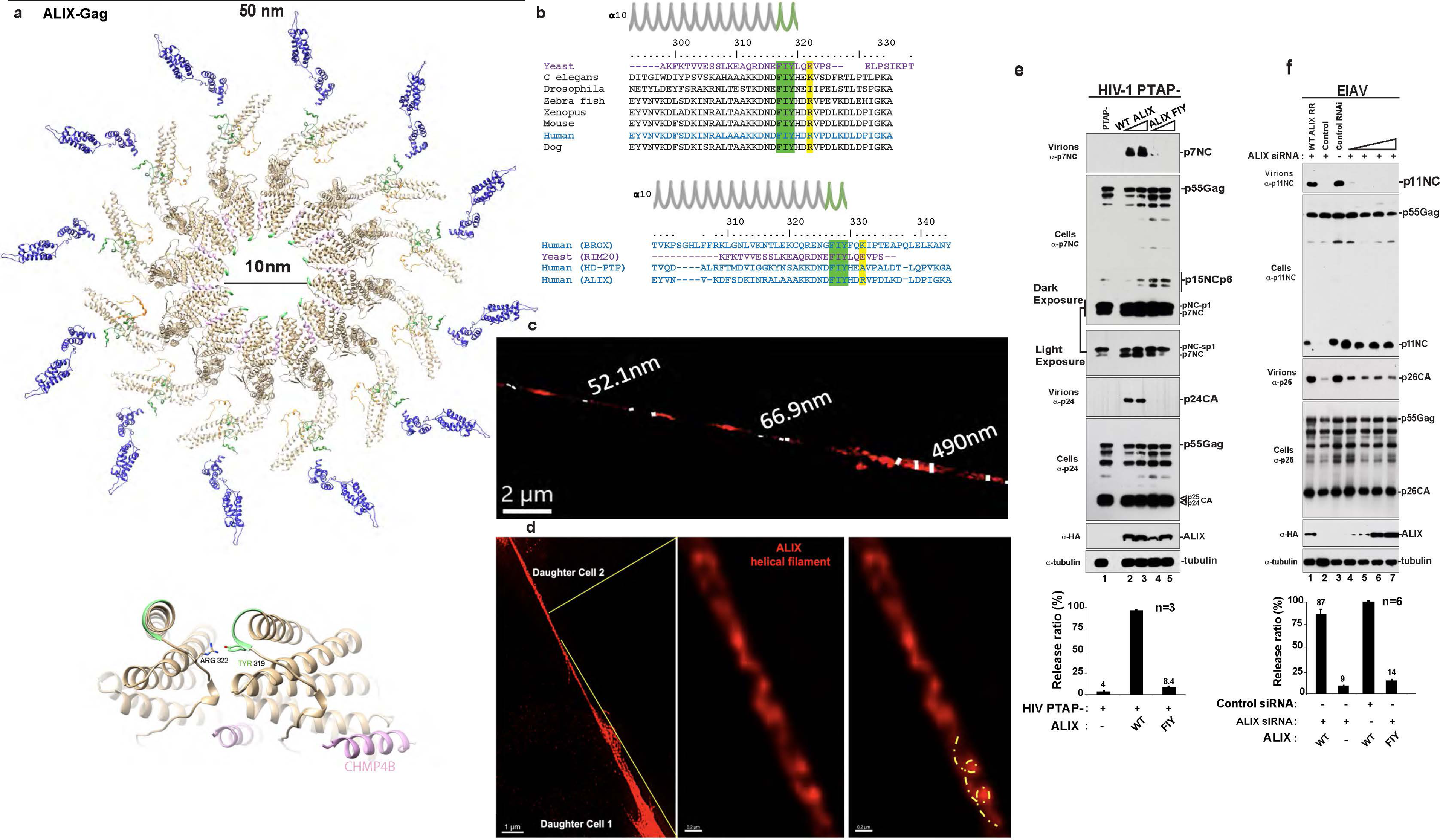
Disruption of a predicted oligomerization interface in ALIX Bro1 domain impairs virus abscission. The FYI motif, an evolutionarily conserved sequence within ALIX is found across several species and Bro1 domain proteins and is involved in dynamic interactions *in crystallo*. (**a)** Molecular model of an idealized HIV-1 budding neck with an ALIX filament recruiting ESCRT-III. Inset zoom shows the position of F319 residue along Alix filament oligomerization interface. (**b**) sequence alignment of the most conserved region of ALIX Bro1 from several model organisms. A second alignment is shown in parallel that reveals complete sequence conservation of the FIY motif from various human and yeast Bro1 domain proteins. Zoom (below alignment) of the site of multimerization within ALIX also reveals the location of the CHMP4B peptide in yellow, between Bro1 domains. The FIY motif is highlighted in pink. (**c**) Super-resolution imaging was used to capture ALIX helical filaments in HeLa living cells. Expression of an ALIX-mcherry fusion protein was used to track ALIX helical scaffold assembly, dynamic membrane tightening and scission within cytokinesis abscission bridges *in vivo*. Alix helical scaffolds were observed within the bridge, and clear helical turns were highlighted with discontinued yellow line. (**d**) Imaris measurement tool was used and estimated helical filament diameter to be ranging from ∼52 nm to 490 nm. (**e**) To validate the nature of the molecular models described above, the FIY motif in ALIX was mutated to alanines and the effect on virus release was assessed. ALIX FIY mutant failed to stimulate the release of an HIV-1 PTAP-mutant (lanes 4 and 5) and processing defective p15NCp1p6 and p9NCp1 accumulated, conversely to WT ALIX that robustly released virus as determined by detection of mature CA and NC released virions (lanes 2 and 3). Data shown are representative of three independent experiments as evaluation below western blot confirms. (**f**) The ALIX FIY mutant was also tested with EIAV, a virus that naturally utilizes ALIX for release. Endogenous ALIX was first depleted by RNAi before WT or FIY mutant ALIX was provided *in trans*. Release of progeny virions and virus maturation were as impaired as HIV-1 PTAP-when ALIX FIY but not WT ALIX was co-expressed. Data shown are representative of six independent experiments as evaluation below western blots recapitulates.

To structurally orient this Bro1 interface for virion scaffolding, we used the parameters of our previous ESCRT-I polymer as well as known structures of ALIX and this newly discovered Bro1 interface to construct a model for an ALIX-stabilized HIV budding neck (**Figure 4a**). We next investigated if ALIX polymers were formed at cytokinetic abscission sites like we previously demonstrated for ESCRT-I filaments. Using super resolution imaging we successfully visualized native ALIX filaments within cytokinesis abscission bridges between two dividing daughter cells. Expression of an mCherry ALIX fusion protein in dividing HeLa cells revealed ALIX helical filaments within abscission bridges and showed clear helices coating inward along the membranous bridges on each side of the midbody furrow site of membrane scission. A representative example of assembled ALIX helical filaments is shown in **Figure 4d**, revealing that ALIX assembles *in vivo* into helical filaments, in agreement with our model in **Figure 4a**. Measurements of ALIX filaments in cells revealed diameters ranging from ∼52 nm to 700 nm (**Figure 4c, and supplemental Figure 3 a and b)**, sizes that correlate with established ALIX functions within viral budding necks, measuring ∼50 nm-as well as cytokinesis abscission bridges whose size is estimated to measure from ∼ 500 nm to 1 um.

To capture ALIX assemblies at sites of function, a dominant negative (DN) approach was adopted, whereby a non-functional truncated form of ALIX comprised of Bro1-V domain was over-expressed in cells with WT HIV-1, similarly to previously described [48]. ALIX Bro1-V carries the Bro1 FIY polymerization interface and the Gag p6 binding domain in the V domain and is therefore expected to bind both in Gag, and make Bro1-Bro1 contacts, without functioning, as the ALIX PRD domain is missing [59]. Over-expression of ALIX Bro1-V inhibited HIV-1 release in a dose-dependent manner [45] and caused nascent virions to accumulate at the PM, where we observed by electron microscopy (EM) an accumulation of a filament-like structures in the lumen of arrested particles (**Supplemental Figure 4**). Their size mostly matched that of predicted viral scission lumens, ∼50 nm. Conversely, no such structures were observed within the lumen of untreated WT HIV-1 (**Supplemental Figure 4),** suggesting that Bro-1-V incorporated in ALIX filamentous polymers in cells and inhibited their proper function at viral budding sites.

The functional significance of ALIX helical filaments observed in **4d** was assessed by targeting their self-assembly interface along the highly conserved F_319_IY motif (**Figure 4b**). The effect of alanine substitutions of FIY to AAA effect on retroviral abscission was assessed by using virus rescue assays of PTAP-HIV-1 and EIAV release as previously described [45, 60]. In the absence of the PTAP L domain, HIV-1 release is nearly absent but can be rescued and significantly enhanced in a dose-dependent manner with the co-expression of WT ALIX (**Figure 4e)**. In addition to increased release of virus, efficient Gag maturation was also recovered as confirmed by the decrease of unprocessed Gag intermediates and NC/NC-sp1 doublet ratio (**Figure 4e**, **lanes 2 and 3-** see light exposure inset). In stark contrast, the FIY/AAA ALIX mutant failed to release HIV-1 (**Figure 4e, lanes 4 and 5**) and EIAV (**Figure 4f, lanes 4-7**) or repair maturation defects, likely due to this mutant’s inability to maintain structural stability of nascent virions, or properly catalyze membrane scission (**Figure 4f, compare lanes 3 to 4-7**). These results show that the ALIX FIY interface is critical for ALIX-dependent inverse topology membrane scission at viral sites. Collectively, the data predicting Bro1-Bro1 as a self-association interface via the FIY motif, combined with ALIX helical models (**Figure 4a**) and ALIX cork-screw filaments observed by super-resolution *in vivo* (**Figure 4d**) support ALIX filaments as structural scaffolds within membranous stalks during inverse topology membrane abscission.

### Both ESCRT-I and ALIX filaments cluster an RNA binding interface critical for inverse membrane scission

Figure 2 showed that virus assembly and budding in the presence of viral RNA genomes robustly enhanced membrane abscission and virus production, suggesting that ESCRT-I as well as ALIX might encounter RNA during membrane scission. To elucidate the underlying mechanisms involved, we probed for interaction(s) between NC, which incorporates viral RNA genomes within developing virions- and ESCRT-I or its subunits. WT NC, WT NC-p1-p6 and their PTAP-mutant counterpart were used in GST-capture assays (**Figure 5**). All Flag-tagged ESCRT-I components were captured with WT NC-p6 (**Figure 5a, lane 3**) while only VPS28, and to a lesser extent, VPS37 and MVB12, were captured with NC-p6 PTAP-mutant or NC alone (**lanes 2 and 4**), indicating there are interactions between these three ESCRT-I components and NC, independently of TSG101 and PTAP. Testing MVB12, VPS37 (**Supplemental Figure 5**) and VPS28 subunits individually (**Figure 5b**) identified NC interactions with VPS28 comparable with both NC alone, and NC-p6 PTAP-mutant. The N-terminal domain (NTD) of VPS28 was found to be both necessary and sufficient for binding NC since interactions between NC and VPS28 NTD were as strong as the full length VPS28, in contrast to the C-terminal domain (CTD) (**Supplemental Figure 5**).

**Figure 5:**
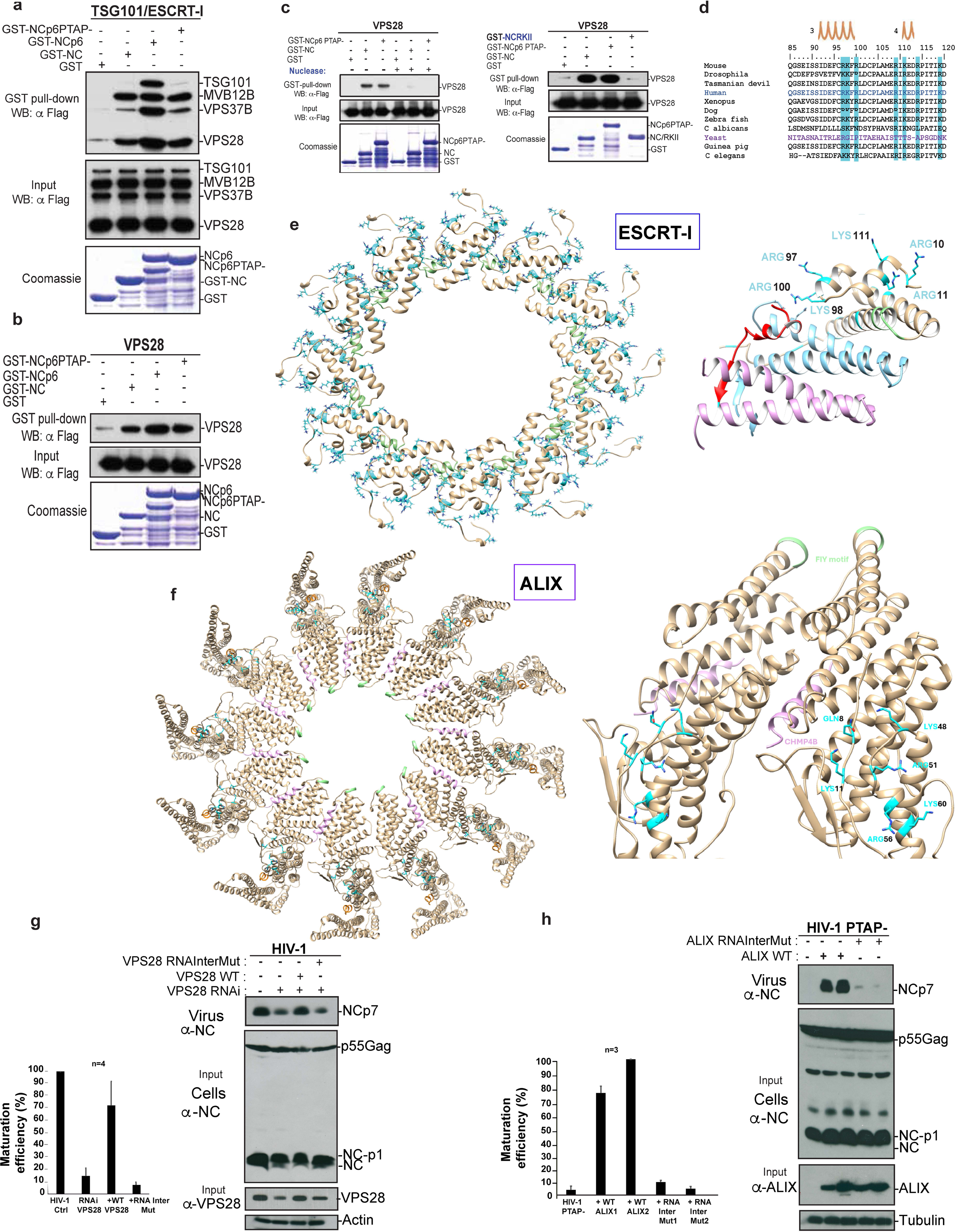
ESCRT-I and ALIX helical filament cluster positively charged interfaces important for scaffolding function: ESCRT-I associates with viral Nucleocapsid (NC) via VPS28 in an RNA-dependent manner. (**a**) GST (lane 1), GST-NC (lane 2), GST-NCp6 (lane 3) or the GST-NC-p6PTAP-mutant (lane 4) fusion proteins expressed in E. coli and captured on glutathione beads were incubated with lysates from 293T cells expressing Flag-tagged WT ESCRT-I subunits TSG101, VPS28, VPS37B and MVB12B (lane 3) or in absence of TSG101 (lane 2). Identical experiments were conducted in parallel with ESCRT-I individual subunits VPS28 (**b**). Additional pulldown assays were conducted between GST-fusion proteins expressing NC, NC-p6PTAP- and the mutant that lack the ability to bind the viral genome devoid NC mutant RKII (**c** left) then followed by incubation with or without benzonase/nuclease (**c** right). GST-fusion proteins purified by glutathione beads were incubated with cell lysates from 293T cells expressing the indicated Flag-tagged VPS28 proteins. In all experiments, captured proteins and cell lysates were analyzed by SDS-PAGE and Western blotting. GST fusion proteins were visualized by Coomassie blue staining. (**d**) The NTD of VPS28 has a positively charged highly conserved surface that contains residues accessible for binding to NC via RNA (blue highlight). Positive residues on VPS28 from the human crystal structure shows that this positive surface is located on the ESCRT-I-viral genome interface (shown in **e left**) and not the ESCRT-I assembly site. ESCRT-I filament formation clusters these positive residues that may promote stronger electrostatic interactions with NC via the RNA genome (shown in **e right**). (**f, left)** ALIX filament formation clusters positive residues in the Bro1 domain that are available to promote electrostatic interactions with NC via the RNA genome. ALIX helical filament formation clusters these positive residues that may promote stronger electrostatic interactions with NC via the RNA genome (shown in **f** right) (**g)** Assessing the role of VPS28 positively charged interface in ESCRT-I mediated HIV-1 release. Virus rescue assays were performed, and HIV-1 abscission recovery was tested with either WT or RNA Interface (InterMut) mutant VPS28 as indicated above the panel. Virion production and Gag expression levels were analyzed by Western blot with an anti-NC antibody. Actin or VPS28 were detected with a mouse anti-Actin and a rabbit anti-VPS28 antibody. The experiment shown is a representative of four independent experiments-value release efficiency graph is shown to the left of the blot. **(h)** To assess the RNA interface functional significance, positively charged residues clustered within ALIX helical filaments were mutated to alanines and the effect on virus release was assessed. ALIX RNAInter mutant failed to stimulate the release of an HIV-1 PTAP-mutant (lanes 4 and 5), conversely to WT ALIX that robustly released virus as determined by detection of mature CA and NC released virions (lanes 2 and 3). Data shown are representative of three independent experiments as evaluation below western blot confirms.

We next asked whether NC association with ESCRT-I VPS28 is RNA dependent was addressed in **Figure 5c**. The NC RKII mutant, which lost the ability to incorporate viral gRNA [61] failed to associate with VPS28 (**5c left**) and a near identical outcome was observed when NC-VPS28 complex was treated with RNAse (**5c right**). Accordingly, a positively charged residue-rich interface forms at the top of TSG101-VPS28 subcomplex helical filaments (Figure **5e**, light blue), where RNA genomes are expected to position within Gag assemblies (**Figure 5e**). These positively charged residues are also highly conserved across species (**Figure 5d**), supporting a role for ESCRT-I in binding nucleic acid. Mutation of residues of this RNA binding interface abrogated ESCRT-I function in HIV-1 budding and abscission from cells **(Figure 5g)**, in line with a model in which ESCRT-I VPS28 RNA binding interface, that clusters only when helical filaments are fully assembled, is important for scaffolding the RNA genome within nascent assembled and budding virions at the PM, while promoting inverse topology membrane scission.

We also found that a patch of positively charged residues clusters in ALIX helical filaments (**Figure 5f)** and that these residues are exposed in the ALIX helical filament model (**Figure 5f, right)**. We hypothesize that this interface is available for interactions with viral genomes, when nascent virions are poised to recruit ESCRT-III within abscission lumens. Disruption of these positively charged surface residues clustered within ALIX polymer (RNAInter mutant) abrogated its function in HIV-1 budding (**Figure 5h**), without affecting virus Gag proteins or ALIX expression in cells, confirming the functional importance of this interface within ALIX helical filaments.

### Assembly of ESCRT-I and ALIX helical filaments is essential for nuclear compartmentalization

Both ALIX and ESCRT-I are key players in cytokinesis completion and rapid abscission of bridges between daughter cells [19, 62, 63]. We showed that both ESCRT-I and ALIX assemble into helical filaments within cytokinesis bridges in living cells (**Figures 3 and 4**). Using this model, the role of filament assembly and their clustered RNA interface was assessed in cytokinesis bridge abscission. Stable cell lines expressing GFP-tubulin and H2B-mCherry were established, to fluorescently track chromosomal segregation during cytokinesis. Cytokinesis delays were monitored following RNAi knockdown of both ALIX and ESCRT-I VPS28 and replenishment with either WT or mutants where FIY and EKYK motifs, along filament assembly interfaces, were abrogated. Significant delays in daughter cells abscission were observed ranging from 20 to 50 min (**Figure 6, compare panels a and c and movie in Supplemental Figure 6)** in RNAi co-depleted cells (Arrested). Reintroduction of WT proteins into RNAi co-depleted cells (Arrested) rescued the phenotype, while ALIX FIY and EKYK VPS28 mutants (Mut Rescue) did not. These results show that ESCRT-I and ALIX require their respective helical filaments oligomerization interfaces in order to assemble and process a timely and efficient inverse topology membrane abscission of cytokinetic bridges. Importantly RNAi co-depletion of ALIX and VPS28 caused additional nuclear defects (**Figure 6c**).

**Figure 6:**
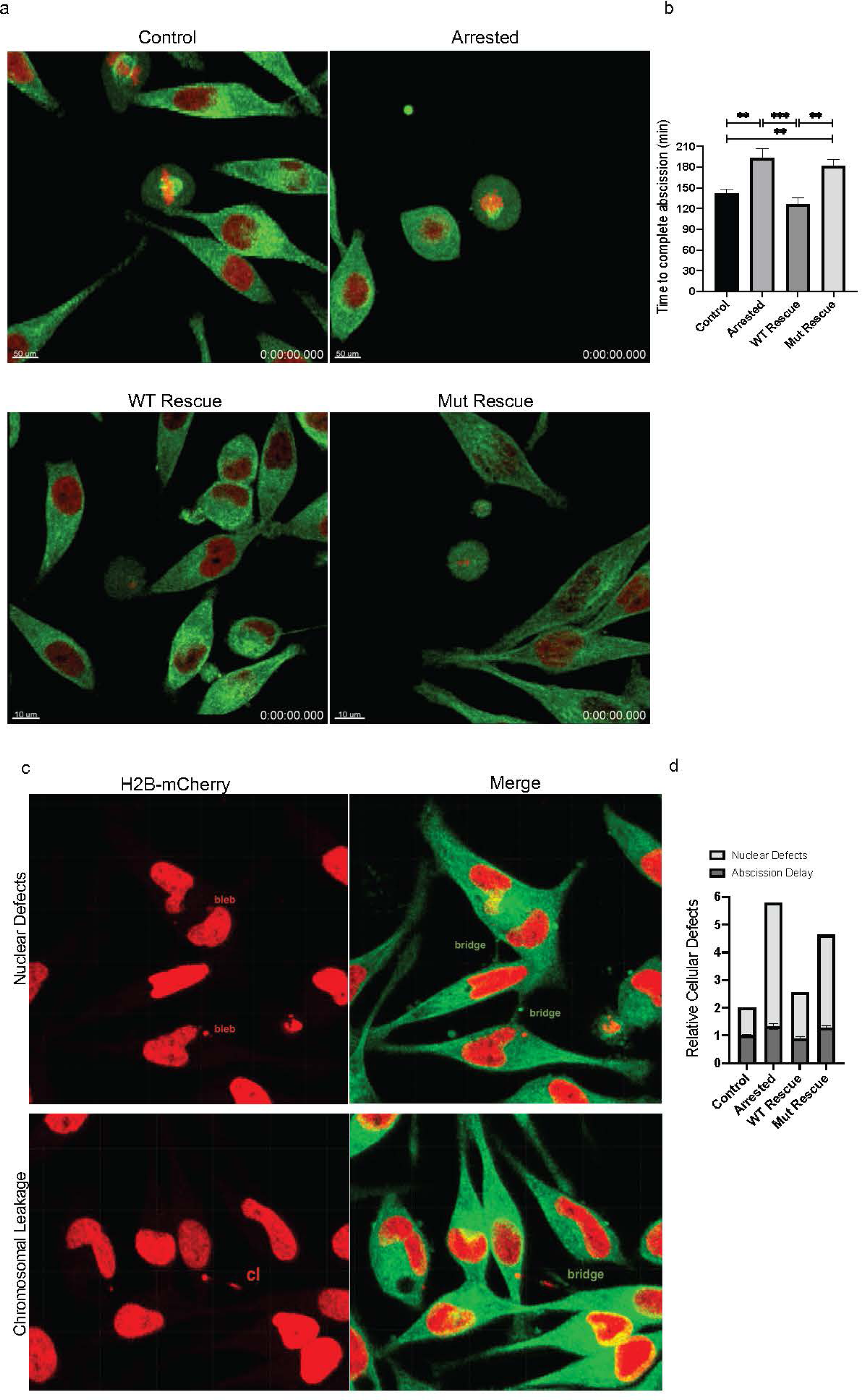
Disruption of ESCRT-I and ALIX helical assemblies impairs cytokinesis and chromosomal segregation. (**a**) HeLa cells constitutively expressing DsRed-H2B and GFP-tubulin were mock transfected (control), co-transfected with RNAi oligos against ALIX and VPS28 alone (Arrested) or in presence of RR expression vectors for either WT ALIX and VPS28 (WT Rescue) or mutants FIY ALIX and EKYK VPS28 (Mut Rescue). Cells were live imaged for 16 hours using a temperature and CO2 controlled Leica SP8 inverted confocal microscope. Knockdown of ALIX and VPS28 by RNAi or Knockdown and reconstitution with FIY and EKYK mutants induced severe delays in cytokinetic abscission by up to 50 minutes compared to the control. Reconstitution with WT variants not only restored, but slightly enhanced abscission completion time (error bars indicate SEM; *n* =300 cells from at least two independent experiments; unpaired *t* test; ***, P < 0.001) (see graphic for evaluations of defects in **b**) Also see Supplemental Figure 5 movie. (**c**) Representative images of “Arrested” phenotypes due to Knockdown or Knockdown and reconstitution of WT rescue or Mut rescue compared to Control cells. “bleb” marks the location of NE breach, “bridge” mark chromosomal genome leak, marked “cl”. NE breaches/defects are marked with arrows in Supplemental Figure 6a. Also see movie in supplemental Figure 6b (**d**) Graphical representation of the defects observed from live imaging studies as described in (**a**) (n>300 cells from at least two independent experiments to determine relative probabilities).

Aberrant nuclei fragmentation (blebbing) and deformations of the nuclear envelope (NE), reminiscent of so-called nuclear herniation [64, 65] were observed in ALIX and VPS28 RNAi co-depleted cells (Arrested) and those reconstituted with mutant ALIX and ESCRT-I VPS28 oligomerization defective mutant proteins (Mut Rescued) (**Figure 6c**). Some of these circular nuclear blebs shed free in the cytoplasm and presented a morphology consistent with chromosomal fragmentation or chromosome failure to properly segregate in daughter nuclei, leading to the formation of micronuclei or micronuclei-like structures (**see 6c and arrows in Supplemental Figure 7a**). Chromosomal leakage between dividing cells was also observed (**6c and Movie in Supplemental 7b**), as evidenced by the presence of H2B-mCherry labelled chromosomes within delayed cytokinetic bridges. In addition, nuclear duplication was also observed, and all nuclei leakage defects were enumerated in **Figure 6d**. Four to five-fold increase in these nuclear defects were observed in both “Arrested” and “Mut Rescued” cells, in comparison to the untreated control or the cells rescued with the WT ALIX and VPS28 proteins (**Figure 6d**). Collectively, this data show that assemblies of ESCRT-I and ALIX into helical filaments and clustering of positive residues promotes temporal control over cytokinesis and reveal their important function in scaffolding and proper segregation of chromosomal nucleic acid within nascent nuclei of dividing daughter cells.

### ALIX and ESCRT-I helical filaments template for ESCRT-III filaments and function synergistically

ALIX or ESCRT-I are recruited to various viral (HIV-1 [15] [8, 16]), and cellular membrane abscission sites including cytokinetic bridges [19], multivesicular bodies [20], and sites of plasma or endosomal membrane repair [21, 22], where they are required for recruitment of ESCRT-III and function in inverse topology membrane scission. Mechanisms of how ALIX and ESCRT-I carry out these roles are incompletely understood. Findings that ALIX and ESCRT-I assemble into helical filaments within cytokinesis bridges **(Figures 3 and 4)** and coat their membranes inward before tightening and processing inverse membrane scission, suggest they might serve as templates for ESCRT-III filaments [29]. To directly test this hypothesis, ESCRT-III was tracked in cytokinesis bridges using CHMP4B-GFP along with ALIX-mcherry, which allowed visualization of ALIX helical filaments therein. Super-resolution imaging of these abscission bridges also revealed CHMP4B filaments that inwardly decorated the cytokinesis abscission bridges and exhibited corkscrew-like pattern which matched the cone-shaped helical pattern displayed by ALIX filaments (**Figure 7a**). Similar results were observed with CHMP4-GFP labeled filaments expressed with ESCRT-I VPS28 filaments assembled along cytokinesis bridges (**Figure 7b**). ESCRT-I VPS28-GFP and ALIX-mCherry co-expression in human cells also showed that both ESCRT-I and ALIX helical filaments co-assembled and were intertwined inwardly along abscission bridges between two dividing daughter cells (**Figure 7c**), indicating synergistical function in living cells. Collectively this data show that helical filaments of ESCRT-I and ALIX are interwoven with ESCRT-III corkscrew like filaments and assemble together along membrane scission stalks in cells, supporting a model where the upstream acting helical filament can initiate ESCRT-III inward recruitment and scaffolding at the membrane, potentially acting as a template for the assembly of its helical polymers, either individually or synergistically.

**Figure 7:**
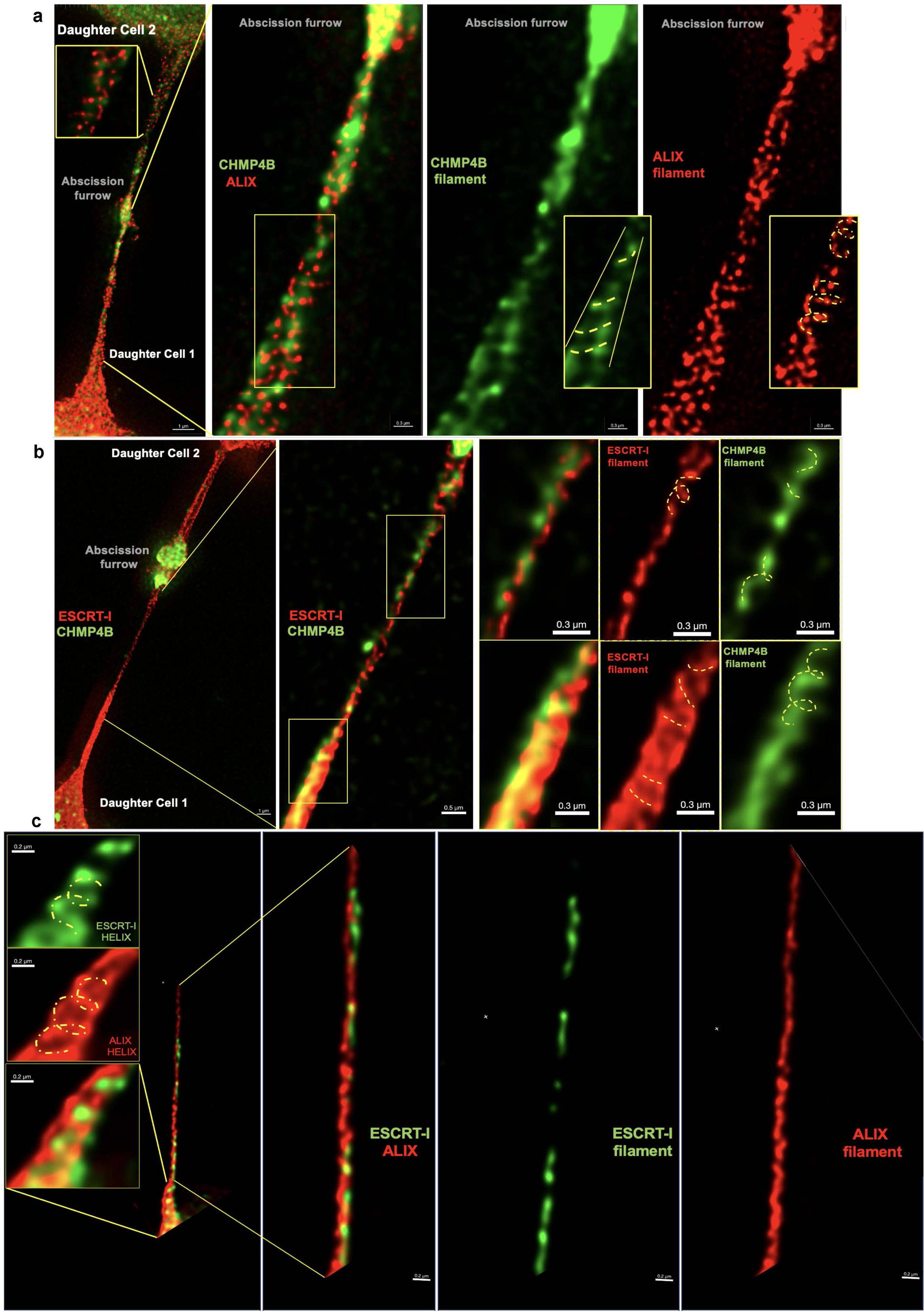
Visualization of helical ESCRT-I, ALIX and ESCRT-III scaffolds function *in vivo*, Representative images from STED imaging of living cells expressing (**a**) ALIX-mCherry and CHMP4B-GFP, (**b**) VPS28-mCherry and CHMP4B-GFP or (**c**) VPS28-GFP and ALIX-mCherry (**c**). Helical turns within membranous bridges are highlighted with discontinued lines. HeLa cells were fixed 20h post-transfection imaging was performed as indicated in Material and Methods. Scale bar sizes are indicated on each image.

## Discussion

While the role of ESCRT-I and ALIX in inverse membrane scission of cellular membrane has been well established [66], the underlying mechanisms involved have remained elusive. ESCRT-I and/or ALIX binding is strictly required for the recruitment and activity of ESCRT-III filaments known to assemble inwardly within inverse topology membrane necks in living cells [29] to complete abscission. Here we show that both ALIX and ESCRT-I assemble into helical filaments in living cells and demonstrate that disruption of filament assembly at highly conserved interfaces across species from yeast to humans interrupted scission of viral and cellular membranes. Remarkably, ESCRT-III CHMP4B filaments inwardly assembled corkscrew-like filaments that matched ESCRT-I and ALIX helical filaments along cone-like membranous abscission bridges, indicating the latter coul template for the assembly of ESCRT-III filaments *in vivo*. Assembled ESCRT-I and ALIX filaments cluster nucleic acid interfaces critical for the proper segregation of viral and chromosomal genomes within the lumens of membrane necks undergoing inverse topology membrane scission. Both ESCRT-I and ALIX filaments also co-assembled within abscission necks indicating synergistical function(s). The most effective scaffolding of cellular (cytokinesis) and viral (budding virions) macrostructures, involving the inward sealing and cutting of enveloping membranes during ESCRT-mediated inverse topology scission, occurs exclusively when ESCRT-I and ALIX filaments act together.

### ESCRT-I and ALIX assembly into helical filaments is essential for inverse topology membrane scission in living cells

Preventing recruitment of ESCRT-I or ALIX to viral budding sites resulted in two defects in addition to membrane scission arrest. These are i) discrete maturation defects illustrated by the gradual appearance and accumulation of uncleaved products of the main structural protein Gag, and ii) loss of an entire lattice of nascent particles that showed thinner shells by electron microscopy. Identical structures were previously seen in lymphocytes, where arrested viruses failed to recruit the ESCRT-I machinery, as well as in the HIV-1 infected T cells MT4 [36] and were described as metastable products of an intermediate maturation stage of developing virions. Our data reveal that both ESCRT-I and ALIX indeed form supportive helical filaments in living cells that assembles inwardly within abscission membranous bridges *in vivo*. Restoring their function repaired maturation defects and provided scaffolding activity within developing virions (Figures 1 and 2). The helical turns of filament sizes were directly observed to reach diameters of up to 500 nm *in vivo*, aligning with the predicted size they occupy within previously observed cytokinesis abscission bridges [54]. We generated structural models of ESCRT-I and ALIX filaments that matched the approximately 50 nm diameter within assembled nascent virions, a size and morphology consistent with observations within viral budding necks and constricting cytokinesis abscission bridges in human cells (**Figures 3, 4 and supplemental Figure 3**). Additionally, our structural models helped define highly conserved helical filament oligomerization interfaces across species and predicted how they encounter viral structural assemblies. Importantly the high conservation of these characteristics, along with that of oligomerization interfaces engaged in both viral and cellular functions support a universal role in which ESCRT-I and ALIX filaments act as early adaptors that may template for ESCRT-III cone-shaped filaments that assemble and inwardly coat membranes (**Figure 7**) to constrict and cleave them in various cellular processes pertinent to health and disease.

### Combined ALIX and ESCRT-I filaments offer the most efficient scaffolding of enveloped macrostructures at the membrane

Early acting ESCRT-I and ALIX are both recruited simultaneously to several viral and cellular membranes. While this dual recruitment initially was interpreted as redundancy, early acting ESCRT-I and ALIX (or a Bro1-domain containing ortholog) were found to be necessary to process inverse topology membrane scission, including at sites of neuronal pruning in Drosophila melanogaster [23], and cytokinesis abscission [24]. Both were also required *in vivo* for the recruitment of ESCRT-III to cytokinetic bridges in Drosophila melanogaster and C. elegans [25], hinting to functional cooperation of both structural scaffolds rather than redundant functions. Indeed, individual depletion of ESCRT-I *or* ALIX caused discrete structural instability of nascent HIV-1 and led to the gradual loss of electron-dense genome-binding lattice from developing virions at the PM (**Figure 1**). Cryo-electron tomograms also captured these defects, and revealed catastrophic loss of viral constituents that comprise the viral protease [3], and the binding domains of the upstream ESCRT components [36, 67]. Consequently, partially processed Gag products accumulated at arrested virions, which displayed thinner shells further confirming structural instability (**Figure 1**). Similarly, infectivity of HIV-1 mutant lacking access to either ESCRT-I or ALIX scaffold was markedly lowered in physiologically significant lymphocytes [45, 61, 68]. These findings highlight the contribution of each upstream ESCRT in providing efficient structural scaffolds to enveloped macrostructures at cellular membranes, cumulatively and cooperatively. Importantly, dual assembly of both ESCRT-I and ALIX helical filaments was observed *in vivo* within abscission bridges (**Figure 7)**, supporting the model in which upstream ESCRT polymers function synergistically to provide the most efficient scaffold at the sites of inverse topology membrane scission, where they also could conceivably template for the assembly of ESCRT-III helical filaments.

### Nucleic acid binding interfaces clustered within ESCRT-I and ALIX filaments are functionally important

The discovery of a new, functionally important, positively charged interface that clusters at the surface of both ESCRT-I and ALIX filaments suggested that they play additional roles along with inverse membrane scission at cellular membranes. High conservation of this charged interface from yeast to human in both ESCRT-I and ALIX filaments emphasized its importance and explains why its disruption interrupted function of both mechanisms of membrane scission (**Figure 5**). Similarly, inhibiting RNA genome binding interface in the Gag structural protein Nucleocapsid (NC) prevented interactions with ESCRT-I VPS28 (**Figure 5**) and ALIX Bro1 domain [69, 70]. The importance of these additional interactions with early ESCRT-I and ALIX were directly illustrated by restoration of membrane scission as well as structural integrity of nascent virions upon supplementation of RNA genome at the plasma membrane (**Figure 2**). These newly uncovered multi-faceted interactions help explain mechanistically why nearly all released enveloped virions contain complete viral RNA genomes (i.e: HIV-1, [71, 72]), underscoring the importance of RNA interaction with upstream ESCRT-I and ALIX in ESCRT-mediated membrane abscission of enveloped viruses. RNA binding interfaces in ESCRT-I and ALIX cluster only in assembled helical filaments, and then become available to interact with nucleic acid of viral or cellular genomes in abscission bridges thus linking upstream ESCRT with the rest of the enveloped virus macrostructure, to which they provide stabilizing structural scaffolding while also processing or repairing membranes by inverse topology membrane scission. In line with these observations, we found that depletion of ESCRT-I and/or ALIX caused leakage of genome associated ribonucleoprotein lattice from nascent virions and inhibited proper chromosomal segregation during cell division. The involvement of additional interfaces that guard against nucleic acid leaks or unstable segregation is conceivable, since ESCRT-II cluster nucleic-acid binding subunits ([73–75]) and ESCRT-III CHMPs are known to localize to cellular or viral membrane gaps where nucleic acid loss or nuclear envelope herniations occur ([76, 77]), in further support of roles in chromosomal segregation via nucleic acid interaction interfaces. These functions also appear to be ancestrally conserved, as archaeal Asgard ESCRT-III subunits were recently found to associate with chromatin [78]. We propose a model in which RNA genomes act as a molecular cement to a multilayered macrostructure, where it binds and assemble viral or cellular proteins on one side of the macrostructure, while engaging the positively charged residues-rich interface clustered on the surface of upstream acting assembling ESCRT-I and helical ALIX filaments on the lower side. These filaments act in turn as scaffolds and support the enveloped macrostructures at the membrane, while simultaneously templating for the downstream acting ESCRT-III filaments to complete membrane envelopment and abscission. The importance of the RNA binding interface in ESCRT-I and ALIX is illustrated with the loss of HIV-1 production upon disruption of its clustered positively charged residues. RNA genome role as a “molecular glue” within enveloped viruses (and likely within other macrostructures or organelles; i.e; nucleus) is consistent with findings that over 90% of HIV-1 released incorporated viral genomes [72], suggesting upstream ESCRT binding through this interface might be part of a selection mechanism that favors the production of virions with the highest infectivity potential. In this study we report that upstream ESCRT-I and ALIX assemble into helical filaments in living cells, and characterise new functional determinants likely key in various membranes sealing pertinent to health and disease, including pathogen production and replication (also reviewed in [79]), cancer [80, 81], neurodegenerative disease [82] and laminopathies [83]. Knowledge derived from molecular mechansims uncovered should help further elucidate processes underlying ESCRT disfunction in human disease and guide the design of novel therapies.

## Material and Methods (see also supplemental material for additional detail)

### Proviral and expression vectors

TSG101 and VPS28 human genes were subcloned into p3XFLAG-(Sigma) pEXPR (IBA) vectors. RNAi Resistant (RR) VPS28 plasmids were constructed by cloning VPS28 modified genes in pcDNA3 (Addgene) to express untagged VPS28 and derivatives. VPS28 mutants were generated based on the structure published in [49] PDB code: 2F6M). Wild-type the (WT) HIV-1 pNL4-3, PTAP−, YPXnL-constructs were previously described in [84]. Provirus EIAVuk is a generous gift from Ronald Montelaro [85].

### Immunoprecipitation assays

Immunoprecipitation assays were conducted as previously described [86].

### Floatation assays

Our flotation assays were initially described by Spearman et al. [33] and modified by Ono and Freed [32] with small modifications (Details are in Supplemental Methods).

### Virus release analysis

293T cells were maintained and transfected and virus released was collected and anylzed as previously described [86].

### RNAi knockdown

293T cells (2.5 × 106 cells/T25) were transfected with 400 pmol or with 100 pmol of Stealth siRNA duplexes against human ESCRT-I genes, individually, or as indicated combinations (Invitrogen). After 36h, cells were cotransfected with the same amount of siRNA duplexes and 1 ug of HIV-1 proviral DNA or the indicated mutant and reconstituted with WT or mutant genes. All knockdowns were assessed by antibody probing for endogenous proteins.

### Pulldown assays and nuclease treatment

HIV-1 NC, NC-p6 or mutant counterparts were expressed in BL21(DE3) pLysS *E. coli* (Stratagene), and their interactions with Flag-tagged ESCRT-I TSG101, VPS28 , VPS37B or MVB12B or mutants, individually or in ESCRT-I, expressed in 293T cells were examined in GST pulldown assays by following the protocol in [86]. **Transmission electron microscopy:** TEM of 293T cells expressing HIV-1 or the indicated mutant was performed as previously described in [61]. Immunogold assays were performed as previously described in [70].

### Cell lines

Stable HeLa cell lines expressing plasmids containing fluorescently tagged GFP-tubulin and histone marker H2b-mCherry (Addgene) were established as described here [87].

### Live Cell Imaging

Cells were imaged using a Leica SP8 inverted confocal microscope equipped with a 63X/1.4NA objective, HyD detectors, 488nm and 561nm lasers. Time lapse movies were typically collected overnight with a time interval of ∼ 4 mins. 4 z-stacks were collected/each time point. At least 30 bridges were analyzed manually to quantify the time to complete abscission.

### Super-resolution imaging and measurements

STED microscopy images were collected with a Leica TCS SP8 STED 3X system (Leica Microsystems) equipped with a HC PL APO 100X/1.4 oil STED white objective, Time-gated HyD detectors, a white light pulsed excitation laser, and 592nm (continuous) and 775nm (pulsed) depletion lasers. STED images were further deconvolved using Huygens Essentials software to reverse optical distortion created during image acquisition. We used the Deconvolution Wizard with automatic background subtraction and microscopic parameters recognition with a continued maximum likelihood estimate (CMLE) iterative algorithm. Filament measurements in 3D were performed using measurement points within the Measurement Ro module of Imaris (version 10.1, Andor Technology Inc., Concord, MA).

### Molecular Modeling

Atomic models for human (hu)ESCRT-I proteins VPS28 and TSG101 were individually constructed in I-TASSER with the yeast ESCRT-I core (PDB code: 2CAZ) as template. Models were then sequentially aligned to the corresponding yeast ESCRT-I using Chimera, which was also used to place monomeric HIV Gag domain p6 on TSG101 (PDB code: 3OBU). ALIX was modeled using the following PDB codes: 2R02, 3UM3, and 4JJY. HIV gag was generated using PDB code: 4USN (CA), PDB code: 2L4L (NC), and PDB code: 2C55 (p6).

## Supporting information

Supplemental Figure 6-linked to Fig6-Movie

## Authors’ contributions

SJS, KMR, SKO, VD and PS co-performed experiments, KMR and PC performed ESCRT-I and ESCRT-I-Gag structural models, VS and ANL conducted proteomics analyses, KN performed TEM imaging and counting, KMR, SY, and SJS performed functional and binding assays, SJS, MS, JK and FB performed STED imagin. FB designed and supervised the project and co-performed experiments, co-analyzed data, and co-wrote the paper.

## Acknowledgments

We thank the AIDS Research and Reference Reagent Program for providing the HIV-1 Gag p24 hybridoma (183-H12-5C) obtained from B. Chesebro, our colleagues Drs. David Ott and Robert Gorelick for the generous gifts of anti-HIV antibodies. This work was supported in part by the Intramural AIDS Targeted Antiviral Program (IATAP) to FB, by the Intramural AIDS Research Fellowship (IARF) to PS and by the NIAID DIR at the NIH to VH, MEG, and SMB.

