## Supplementary figures and images for "Human ESCRT-I and ALIX function as scaffolding helical filaments *in vivo*"

### Supplemental Figure 1

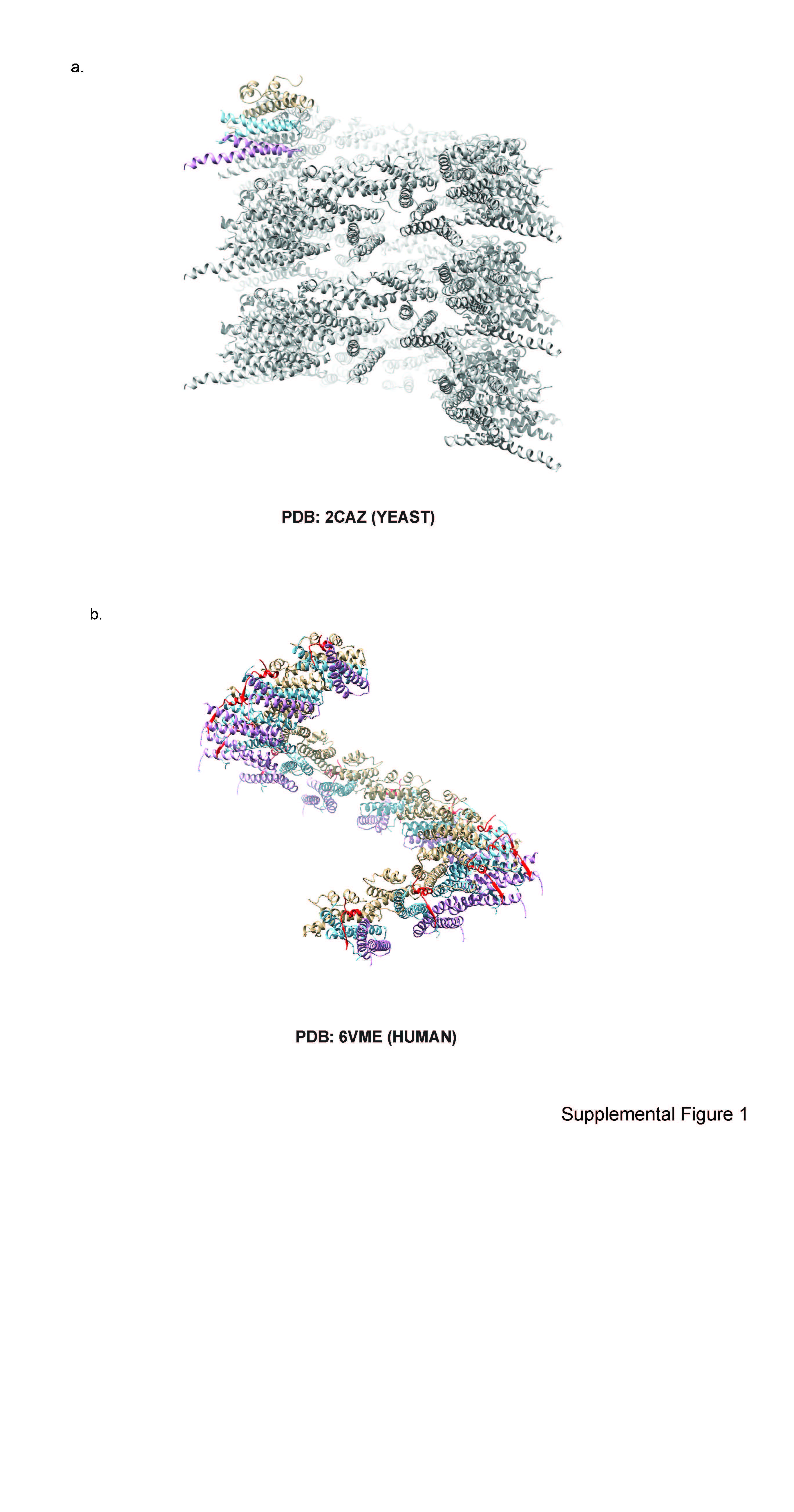

### Supplemental Figure 2

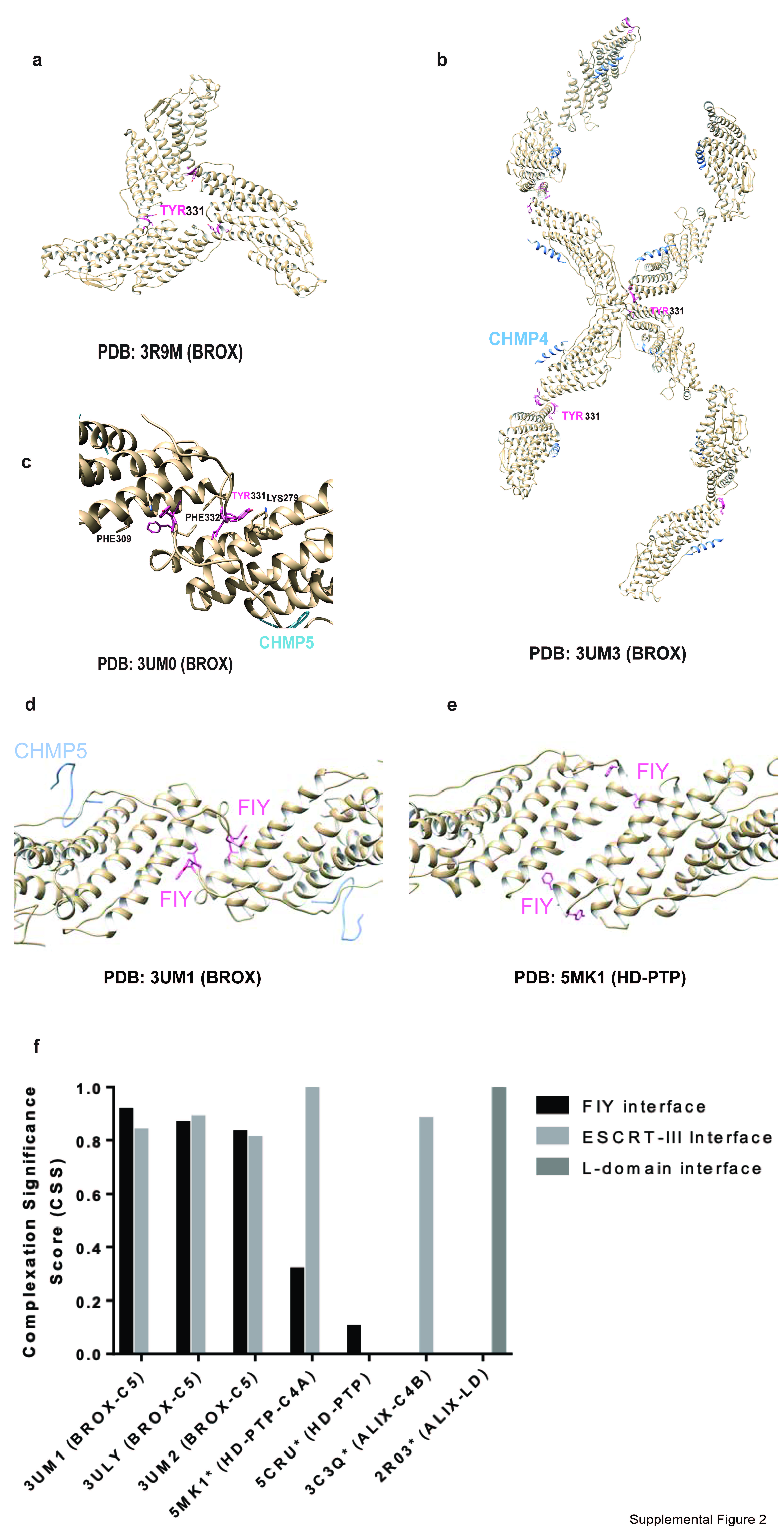

### Supplemental Figure 3

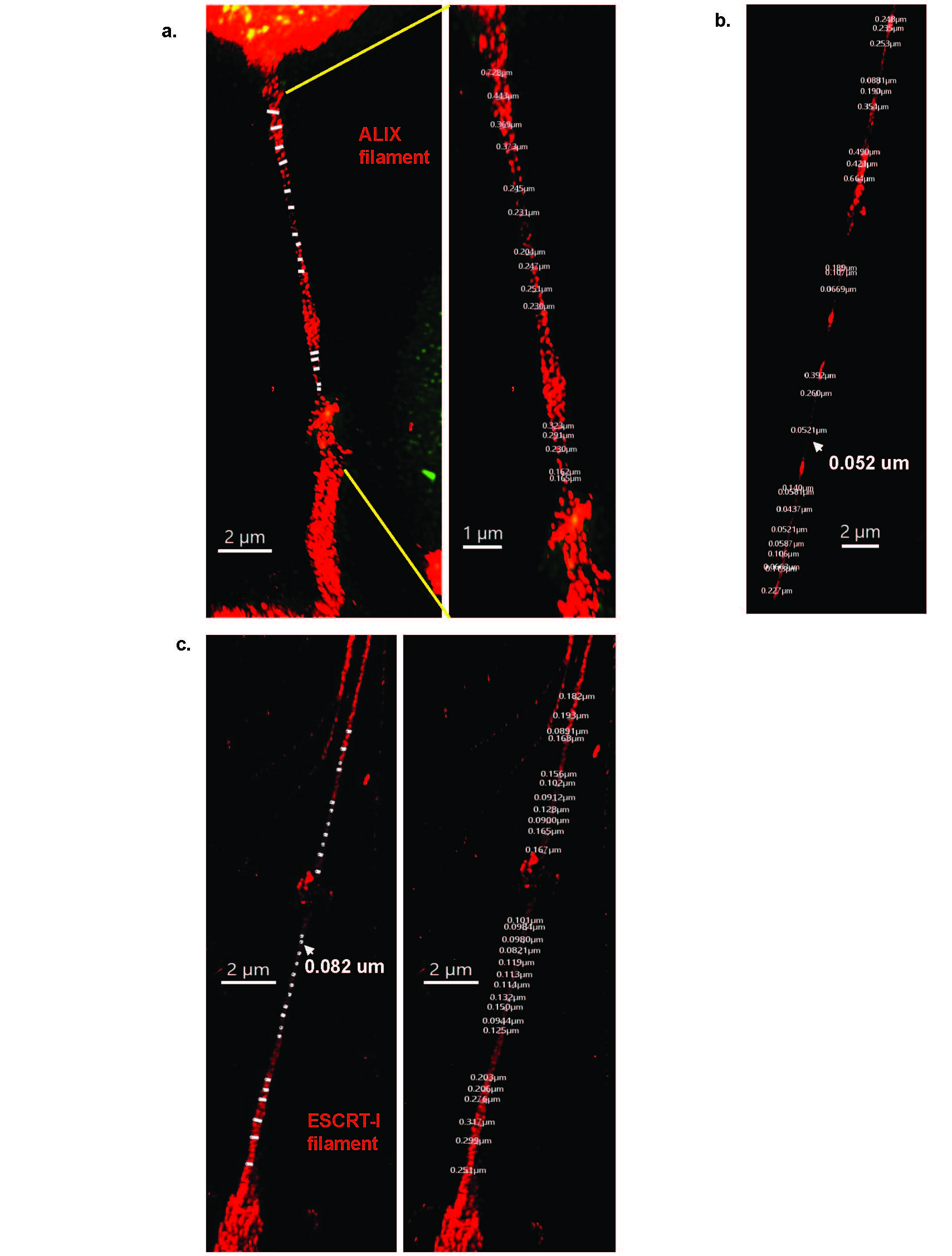

### Supplemental Figure 4

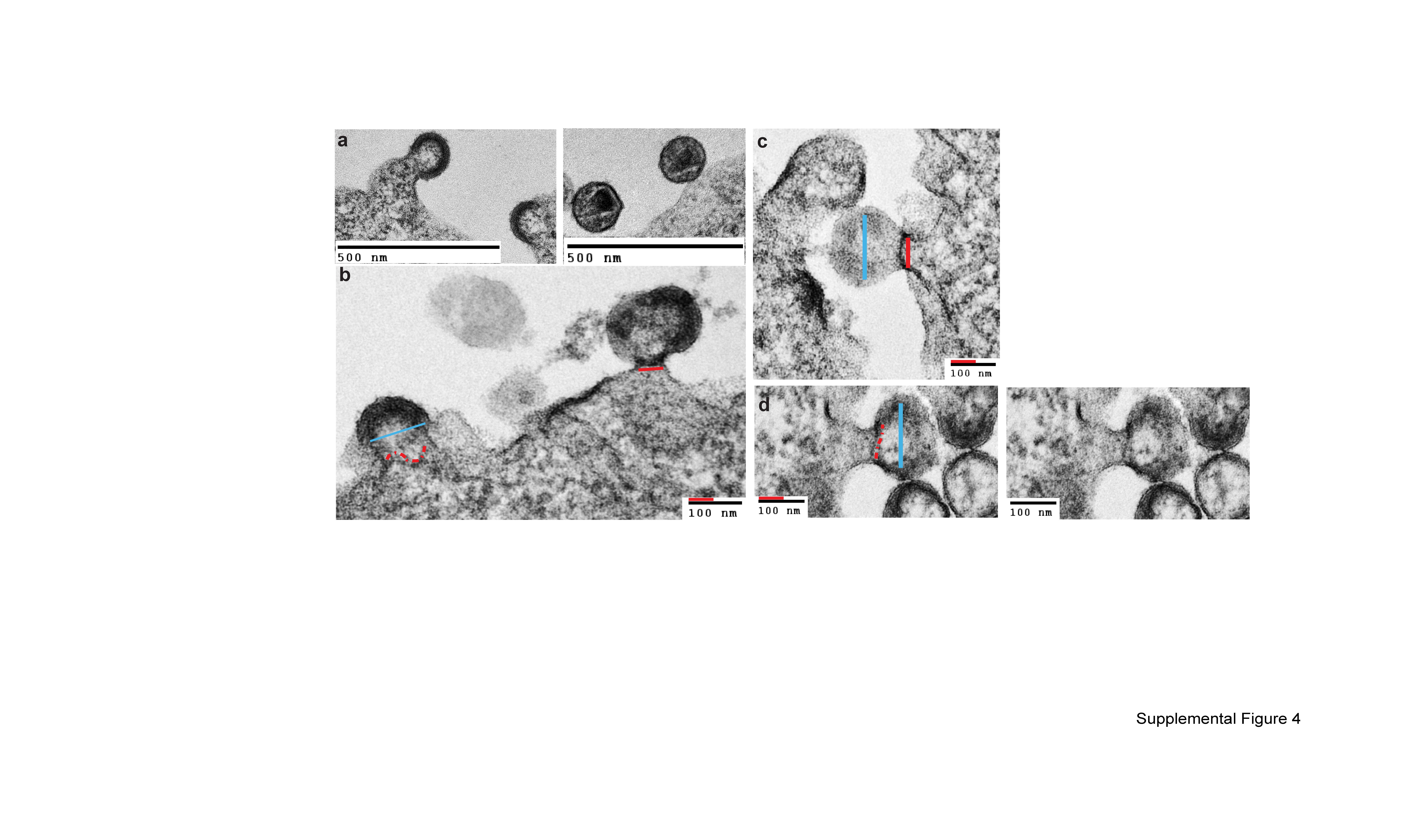

### Supplemental Figure 5

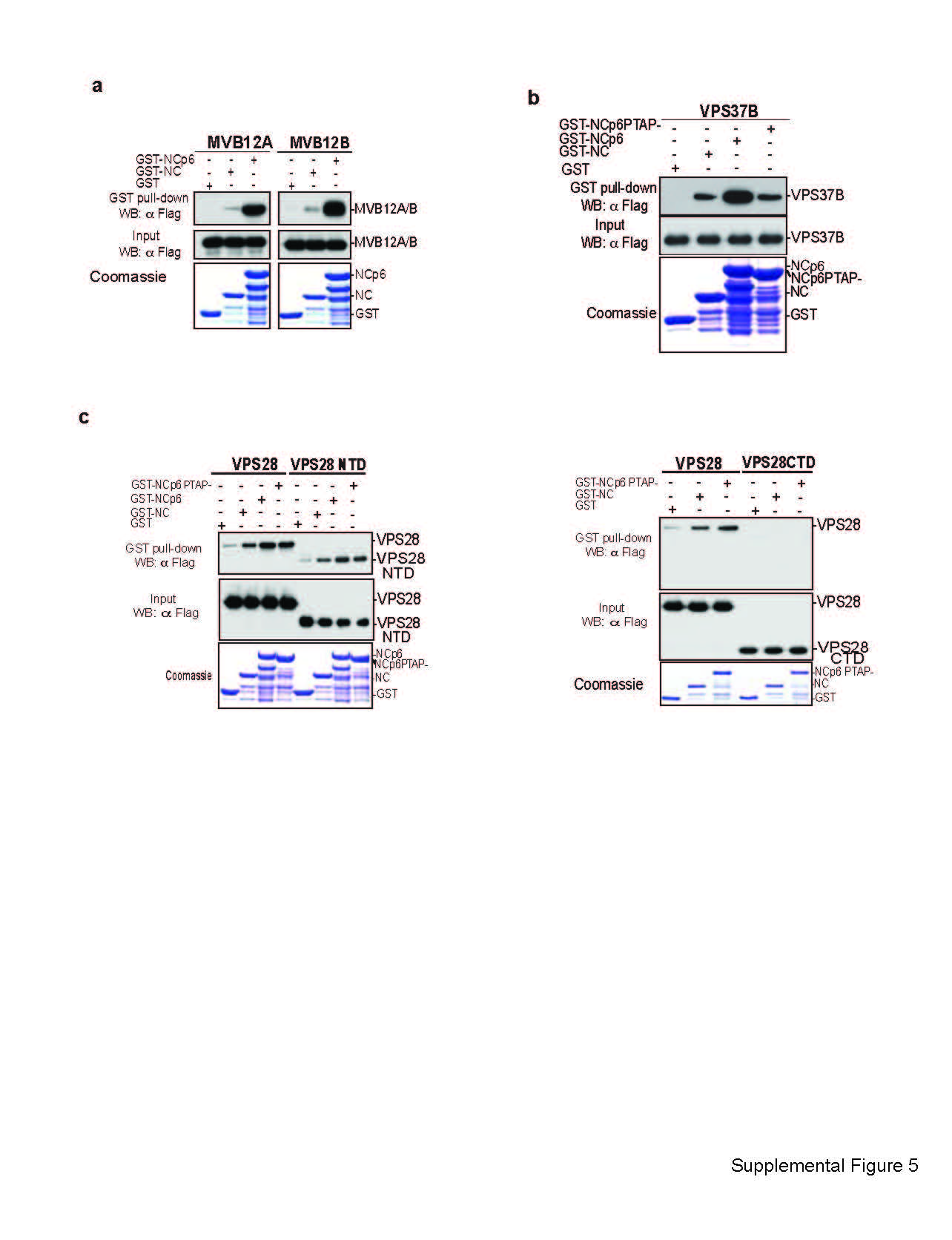

### Supplemental Figure 7a

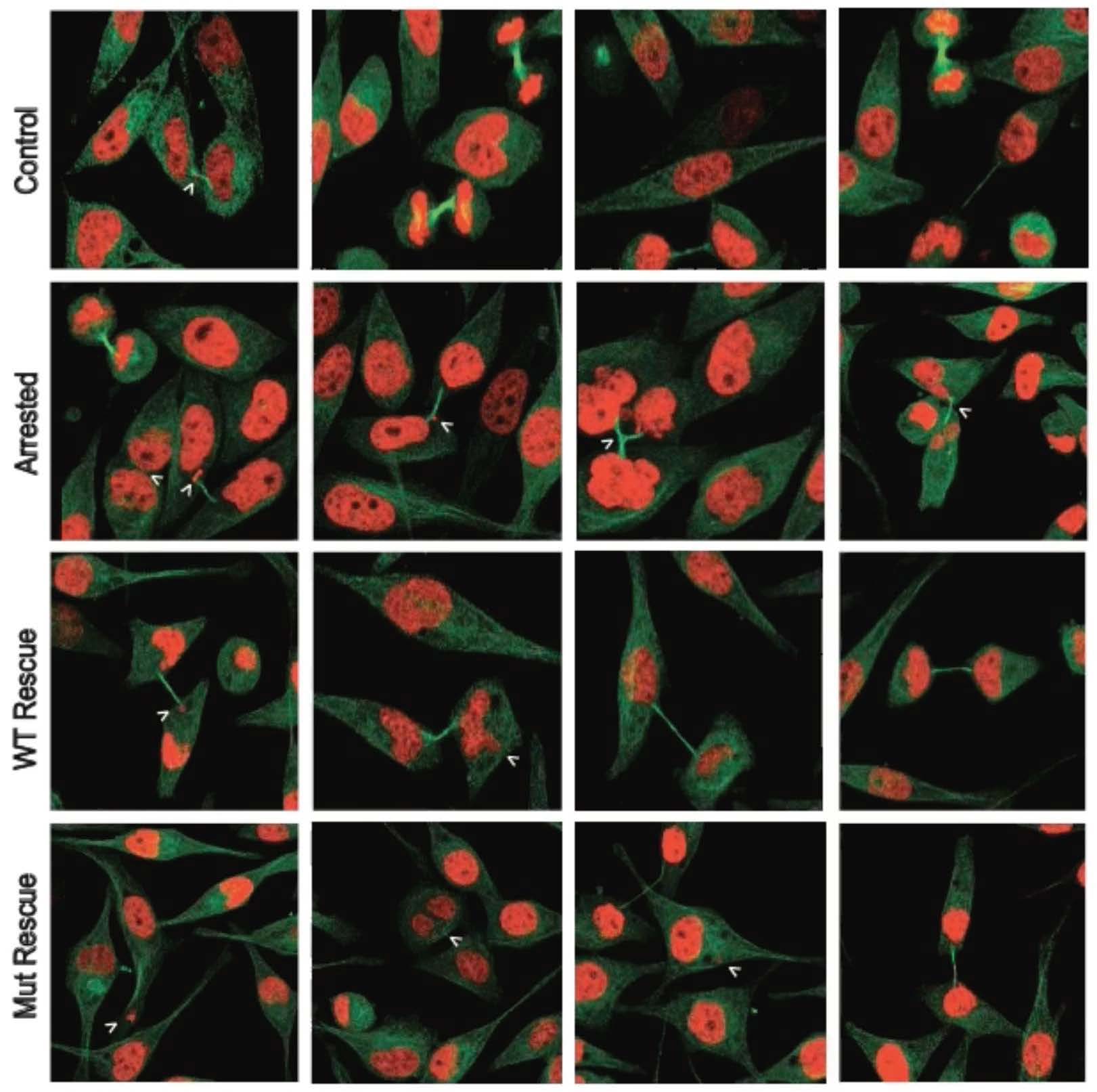
