## Supplemental Figure legends for "Human ESCRT-I and ALIX function as scaffolding helical filaments *in vivo*"

**Supplemental Figure legends: Spada et al. 2024**

**Supplemental Figure 1:** Yeast ESCRT-I forms filaments in crystal structures with an inner diameter of roughly 10 nm (PDB code: 2CAZ). (**b**) Structure of a Human ESCRT-I crystal filament that was resolved during this study (PDB code: 6VME).

**Supplemental Figure 2: Analysis of crystallographic interfaces from ALIX paralogues Brox and HD-PTP. (a)** Brox forms an array of multimers within crystal structures that revolve around FIY motifs (pink). **(b)** ALIX ortholog Brox in complex with an ESCRT-III CHMP4B peptide adopts a spacious intersecting wire crystal packing geometry composed of several tetrameric and dimeric interfaces about the FIY motif in Brox. (**c**) Representative image of a HD-PTP dimer interface from crystal structures that also involve FIY motifs. (**d**) In the presence of CHMP5, Brox crystals pack as dimers with a disordered pair of FIY motifs at the site of interaction. **(e)** A pair of Bro1 dimers are packed together by two FIY motifs in HD-PTP. (**f**) Complexation Significance Scores (CSS) from several Brox and HD-PTP crystal structures reveal that Bro1-Bro1 assemblies about FIY motifs are likely biologically relevant. Three instances of Brox Bro1 domain multimers in complex with CHMP5 (Brox-C5) displayed CSS scores of 0.8 or greater, indicating biological significance on a scale of 0 to 1. In two of these three instances the CSS scores were greater than that of the Brox-ESCRT-III interface. In the case of HD-PTP alone or in complex with CHMP4A (HD-PTP-C4A), FIY interfaces between Bro1 domains received much lower CSS scores, but the crystal packing of these FIY motifs was much more spatially separated than that of Brox. Generally, CSS scores between Bro1 domains positively correlated with FIY motif proximity. In the case of ALIX, no crystallographic Bro1 interfaces using FIY motifs could be found and no other Bro1 oligomers with CSS scores greater than 0 could be identified. As a comparison, ALIX in complex with CHMP4B (ALIX-C4B) or an L-Domain peptide (ALIX-LD) displayed CSS scores comparable to the FIY interfaces found within Brox crystal structures, in further support of a model for FIY-mediated Bro1 oligomerization.

**Supplemental Figure 3:** Filament measurements in 3D were performed using measurement points within the Measurement Ro module of Imaris (version 10.1, And or Technology Inc., Concord, MA). (**a**) and **(b)** ALIX filament occupy a diameter ranging from ~ 52 nm to 1um in representative abscission bridges between two dividing cells. **(c)** An example of ESCRT-I helical filament, within a representative abscission bridge between two dividing cells, which occupy a diameter ranging from ~82 nm to 1um.

**Supplemental Figure 4: Electron micrographs from HIV-1 infected cells compared to infected cells also over-expressing ALIX Bro1-V fragment.** Filamentous electron dense structures at the bases of budding necks were observed (**b,c, and d**) that are likely ALIX assemblies of approximately 50 nm diameter, a size in agreement with that seen with ALIX filaments observed *in vivo* in Figure 4d. They are absent in WT nascent virions are expressed alone (**a**). Blue lines mark nascent virions, while the red lines mark ~ 50 nm sites of filamentous assemblies in budding necks.

**Supplemental Figure 5 (linked to Figure 5 in main text): ESCRT-I subunits MVB12A/B do not bind HIV-1 NC, in contrast to VPS28 NTD.** GST (lane 1), GST-NC (lane 2), GST-NCp6 (lane 3) or GST-NC-p6PTAP- mutant (lane 4) fusion proteins expressed in E. coli and captured on glutathione beads were incubated with lysates from 293T cells expressing Flag-tagged WT ESCRT-I subunits MVB12A, MVB12B (**a**) or VPS37B (**b**). (**a**) MVB12A/B does not bind NC and VPS37 retains some binding to NC. (**c**) VPS28 CTD (117aa-end) does not bind NC in contrast to the VPS28 N-terminal (NTD 1-120aa). In all experiments, captured proteins and cell lysates were analyzed by SDS-PAGE and Western blot using indicated antibodies. GST fusion proteins were visualized by Coomassie blue staining.

**Supplemental Figure 6-Movies linked to Figure 6a in main text: Disruption of ESCRT-I and ALIX helical scaffolds delays cytokinesis** HeLa cells constitutively expressing DsRed-H2B and GFP-tubulin were mock transfected (control), co-transfected with RNAi oligos against ALIX and VPS28 alone (Arrested) or in presence of RNAi resistant expression vectors for either WT ALIX and VPS28 (WT Rescue) or mutants FIY ALIX and EKYK VPS28 (Mut Rescue). Cells were live imaged for 16 hours using a temperature and $CO_2$ controlled Leica SP8 inverted confocal microscope. Knockdown of ALIX and VPS28 by RNAi or Knockdown and reconstitution with FIY and EKYK mutants induced severe delays in cytokinetic abscission by up to 50 minutes compared to the control. Conversely, reconstitution with WT variants not only restored, but slightly enhanced abscission completion time (error bars indicate SEM; $n$ =300 cells from at least two independent experiments; unpaired $t$ test; ***, $P < 0.001$) and quantified in Figure 6**b** in main text.

**Supplemental Figure 7-Movie linked to Figure 6c in main text:** (**a**) "Arrested" phenotypes due to Knockdown or Knockdown and reconstitution of WT rescue or Mut rescue compared to Control cells.. Nuclear envelope breaches or defects are marked with arrows. Quantification of the defects observed from live imaging studies as described in (Supplemental Figure 6) (n>300 cells from at least two independent experiments to determine relative probabilities are shown in Figure 6**d** in main text. (**b**) Live imaging movies showing nuclear defects and chromosomal leaks described in still images in Figure 6c and supplemental Figure 7**a**.
