## Supplemental Material and Methods for "Human ESCRT-I and ALIX function as scaffolding helical filaments *in vivo*"

**Proviral and expression vectors:** Human ESCRT-I subunit genes of TSG101 and VPS28 were subcloned into p3XFLAG-myc-CMV-26 vector (Sigma) to obtain the N-terminally 3XFLAG tagged protein expression plasmids and cloned in pEXPR plasmid (IBA) to generate an expression plasmid for a strep-tagged version of the ESCRT-I subunits. RNAi Resistant (RR) VPS28 expression plasmids were constructed by cloning VPS28 modified appropriate gene in pcDNA3 (Addgene) to express untagged VPS28 and derived proteins. VPS28 M39D/F43D mutant was generated based on the structure published in in reference 49 (PDB code: 2F6M); M39 and F43 residues in the human VPS28 gene were mutated to D. The EKYK VPS28 mutant was generated by substituting residues underlined in the $_{52}$EKAYIKDC$_{61}$ sequence to alanine. The ALIX FIY mutant was generated by substituting all three residues to alanine. We used the wild-type (WT) HIV-1 pNL4-3, the L-domain singly defective HIV-1 mutant PTAP− , YPXnL-; proviral constructs were previously described Dussupt et al, 2009, reference 45. EIAVuk proviral construct is a generous gift from Ronald Montelaro (University of Pittsburgh) and was previously used in Bello et al, 2012, reference 85.

**Floatation assays:** Our flotation assays were initially described by Spearman et al. 1997, reference 33 and modified by Ono and Freed, 1999, reference 32 with small modifications. 293T cells were collected and washed with cold PBS, resuspended in 10 mM Tris-HCl containing 1 mM EDTA, 10% (wt/vol) sucrose, and Complete protease inhibitor cocktail. Postnuclear supernatants (PNS) were obtained after sonication of cell suspensions in ice water and spun at 1000g at 4°C to obtain PNS. Next, 250 µl of PNS was mixed with 1.25 ml of 85.5% (wt/vol) sucrose in TE and placed on the bottom of a centrifuge tube. Seven ml of 65% (wt/vol) sucrose in TE were layerd on top of PNS-containing 73% (wt/vol) sucrose mixture, and finally 3.25 ml of 10% (wt/vol) sucrose in TE was layered at the top of gradients. Gradients were centrifuged at 100,000 × g for 18 h at 4°C in a Beckman SW41 rotor and twelve 1-ml fractions were collected from the top of the centrifuge tube for western blot analysis using anti-MA or anti-NC antibodies. PNS were treated with 1M NaCl or RNAse (Novagen) for 30 min on ice prior to mixing with 85.5% sucrose (bottom of gradient).

**Immunoprecipitation assays:** Immunoprecipitation assays were conducted as previously described in Sette et al, 2016, reference 69. Briefly, 293T cells (2 × 106 cells/T25) were seeded into T25 flasks and transfected the following day using Lipofectamine 2000. At time 48h posttransfection, the cells were lysed in Lysis buffer (1% (v/v) Igepal, 0.1% (w/v) sodium dodecyl sulfate, 0.5% (w/v) Sodium deoxycholate or RIPA buffer (140 mM NaCl, 8 mM Na2HPO4, 2 mM NaH2PO4, 1% Nonidet P40 [NP-40], 0.5% sodium deoxycholate, 0.05% sodium dodecyl sulfate [SDS]) and Complete protease inhibitor cocktail [Roche, Indianapolis, IN]). Immunoprecipitation complexes and cell lysates (input fractions) were analyzed by SDS-PAGE and western blot using the indicated antibodies.

**Virus release analysis:** 293T cells were maintained and transfected as previously described Sette et al, 2016, reference 69. Twenty-four hours after transfection, cells and culture media were harvested and their protein content was analyzed by SDS-PAGE and western blot using the indicated antibodies. HIV-1 proteins were detected using an anti-HIV-1 p24 monoclonal antibody (clone 183-H12-5C) or NEA-9306 (PerkinElmer).

**RNAi knockdown:** 293T cells (2.5 × 106 cells/T25) were transfected with 400 pmol or with 100 pmol of Stealth siRNA duplexes against human TSG101, ALIX, VPS28, or indicated combinations (Invitrogen life technologies). After 36h, cells were cotransfected with the same amount of siRNA duplexes and 1 ug of HIV-1 proviral DNA or the indicated mutant and reconstituted with WT or mutant VPS28 and ALIX construct(s). Cells and virus were harvested and processed as described above. All knockdowns were assessed by western blot of cell extracts and antibody probing for endogenous proteins.

**Pulldown assays and nuclease treatment:** The empty pGEX vector or that carrying the coding sequences of HIV-1 NC, NC-p6 or mutant counterparts were expressed in BL21(DE3) pLysS *E. coli* (Stratagene), and their interactions with Flag-tagged ESCRT-I TSG101, VPS28 , VPS37B or MVB12B and its mutants, individually or in an ESCRT-I complex, expressed in 293T cells were examined in GST pulldown assays by following the protocol previously described Sette et al, 2016, reference 69. Where indicated, protein complexes captured on beads were incubated for 30 min at 37°C in the presence or absence of 75 U (0.75 U/µl) benzonase/nuclease (Novagen) in benzonase buffer (1.2 mM $MgCl_2$, 50 mM Tris-HCl [pH 8.0]). Eluted complexes and cell lysates (input fractions) were analyzed by SDS-PAGE and western blotting using the indicated antibodies.

**Transmission electron microscopy:** TEM of 293T cells expressing HIV-1 or the indicated mutant was performed as previously described in Dussupt et al, 2009, reference 45. Examination and counting of ~ 300 to 400 assembly events at the PM were performed to evaluate budding phenotypes and/or particle morphogenesis of HIV-1 WT or the indicated mutant. Immunogold assays were performed as previously described in Sette et al, 2016 reference 70.

**Establishment of Cell lines for Live Cell Imaging:** Stable HeLa cell lines expressing plasmids containing fluorescently tagged GFP-tubulin and histone marker H2B-mCherry (Addgene) were established as previously described in Thoresen et al, 2014, reference 87. Cell lines were depleted of endogenous proteins and reconstituted with WT or mutant variants as described above.

**Live Cell Imaging:** Cells were plated in poly-lysine coated coverslip bottom chamber (Nunc) and imaged using a Leica SP8 inverted confocal microscope equipped with a 63X/1.4NA objective, HyD detectors, 488nm and 561nm lasers, adaptive focus control, and an environmental chamber with temperature control and CO2. Time lapse movies were typically collected overnight with a time interval of approximately 4 minutes. 4 z-stacks were collected at each time point. At least 30 bridges were analyzed manually per sample to quantify the time to complete abscission.

**Molecular Modeling of Viral and Host Proteins:** Atomic models for the human ESCRT-I proteins VPS28 and TSG101 were individually constructed using the Iterative Threading Assembly refinement method (I-TASSER) with the yeast ESCRT-I core (PDB code: 2CAZ) as the structural template. Human protein models were then sequentially and structurally aligned to the corresponding yeast ESCRT-I proteins using the Matchmaker extension in UCSF Chimera. This extension was also used for the placement of monomeric HIV p6 protein of the full-length HIV Gag protein on the N-terminal domain of TSG101 (PDB code: 3OBU). The resulting model was assessed and refined with MolProbity for the completion of human ESCRT-I filaments. The remaining ESCRT-I components (VPS37 and MVB12) were modeled in last by aligning the yeast heterotetramer core structure to the human filament created here (PDB code: 2P22). In order to model an ALIX multimer in the ~50 nm lumen of HIV scission necks, the ideal parameters identified from the ESCRT-I filament were used as a template. To match the geometry and placement of HIV Gag as determined by our ESCRT-I filament described above, an "extended ALIX" was first modeled where the Bro1 domain and distal portion of the V domain resemble ALIX from known crystal structures (PDB code: 2OEV), but the proximal half of the V domain is an extended conformation as determined by crystallography (PDB code: 4JJY). The full-length Gags described herein were then docked onto this extended ALIX model.

Full-length HIV Gag molecules were constructed entirely from known structures in the Protein Data Bank (rcsb.org) that constitute the various individual domains of the multi-domain protein. Our Gag model begins with the immature Capsid-SP1 lattice 18 derived from viral particles (PDB code: 5L93) that displays a six-fold symmetry. A multimer of HIV NC (PDB code: 1A1T) molecules corresponding with the same six-fold symmetry was constructed using the HOMOMER function of the GalaxyWEB server (galaxy.seoklab.org/) by manually inputting an oligomeric state of 6. In a similar fashion, an HIV p6 hexamer (PDB code: 2C55) was created using the HSYMDOCK Server (huanglab.phys.hust.edu.cn/hsymdock/) and manually assigning a cyclical symmetry of 6. The full-length HIV Gag was then constructed by ab initio free docking of the HIV NC hexamer onto the C-terminus of the HIV CA immature lattice using the HDOCK server (hdock.phys.hust.edu.cn/) and selecting the resulting complex with the lowest docking score, indicative of the complex formed with the lowest energy requirements. This protocol was repeated for the incorporation of the HIV p6 multimer and completion of a full-length immature HIV Gag hexamer.
